# Systematic sequencing of chloroplast transcript termini from *Arabidopsis thaliana* reveals >200 transcription initiation sites and the extensive imprints of RNA-binding proteins and secondary structures

**DOI:** 10.1101/621938

**Authors:** Benoît Castandet, Arnaud Germain, Amber M. Hotto, David B. Stern

## Abstract

Chloroplast transcription requires numerous quality control steps to generate the complex but selective mixture of accumulating RNAs. To gain insight into how this RNA diversity is achieved and regulated, we systematically mapped transcript ends by developing a protocol called Terminome-Seq. Using *Arabidopsis thaliana* as a model, we catalogued >215 primary 5’ ends corresponding to transcription start sites (TSS), as well as 1,628 processed 5’ ends and 1,299 3’ ends. While most termini were found in intergenic regions, numerous abundant termini were also found within coding regions and introns, including several major TSS at unexpected locations. A consistent feature was the clustering of both 5’ and 3’ ends, contrasting with the prevailing description of discrete 5’ termini, suggesting an imprecision of the transcription and/or RNA processing machinery. Numerous termini correlated with the extremities of small RNA footprints or predicted stem-loop structures, in agreement with the model of passive RNA protection. Terminome-Seq was also implemented for *pnp1-1*, a mutant lacking the processing enzyme polynucleotide phosphorylase. Nearly 2,000 termini were altered in *pnp1-1*, revealing a dominant role in shaping the transcriptome. In summary, Terminome-Seq permits precise delineation of the roles and regulation of the many factors involved in organellar transcriptome quality control.

## INTRODUCTION

The sophisticated interplay of factors regulating chloroplast gene expression results from over a billion year symbiosis between the plastid and nucleus. Transcription of the entire plastome (1, 2) combined with inefficient termination (3, 4) leads to a complex primary transcriptome that undergoes numerous maturation steps. These may include 5’ and 3’ end processing, intergenic cleavage of polycistronic transcripts, intron removal through RNA splicing, and RNA editing to convert specific cytosines into uracils (5–7). Among the protein factors involved in maturation, a variety of endo- and exoribonucleases (RNases) are responsible for processing and cleavage (8), while RNA-binding proteins (RBPs) counteract their activities to stabilize some transcript ends (9), or participate in splicing or editing protein complexes.

Legacy chloroplast RNA maturation analyses have used tedious gene by gene molecular techniques, restricting most detailed RNA analyses to several mono- or polycistronic transcripts including *psbA, rbcL, atpI*-*atpH*-*atpF*-*atpA*, and *psbB*-*psbT*-*psbN*-*psbH*-*petB*-*petD* (10), along with the ribosomal RNA operon, which together may not be fully representative of plastid RNA maturation pathways. Gene-by-gene analyses of processing factors may also have limited value. For example, the RNase CSP41a was shown to possess strong endoribonuclease activity *in vitro* (11), however *in vivo* mutant analysis suggests regulatory roles that may have little to do with RNase activity *per se* (12–15). Additionally, while mutants for the endoribonuclease RNase E, its specificity partner RHON1, and the 5’→3’ exoribonuclease and endoribonuclease RNase J accumulate novel and apparently unprocessed transcripts, pleiotropic effects are prone to masking their precise sites of cleavage or interaction (16–18).

To overcome some of these limitations, we and others have increasingly employed RNA-Seq-based approaches that yield genome-wide cataloging and mechanistic insights into chloroplast transcription, editing, splicing, and translation (19–28). Our own results, for example, revealed a large number of non-coding RNAs, which additional research suggests may include a class that exerts its functions through sense-antisense RNA pairing (19, 29, 30). It is difficult to fully understand transcript function, however, without knowledge of the 5’ and 3’ termini which together help define promoter sequences, regulatory UTRs, and the potential for sense-antisense pairing. On a genome-wide scale, we refer to these 5’ and 3’ ends as the (RNA) terminome, and the associated technique as Terminome-Seq.

Efforts to define the chloroplast RNA terminome have been limited to date, with the most comprehensive study focused on 5’ ends in barley, a model that has been effectively used to dissect the respective roles of nucleus- and chloroplast-encoded RNA polymerases (31, 32). On a genome-wide level, barley chloroplasts were found to possess larger than expected numbers of both primary and processed 5’ termini, consistent with a highly complex transcriptional landscape (20). We chose Arabidopsis for our analysis, because of its broad use to dissect post-transcriptional RNA events in the chloroplast, including the analysis of RNase mutants, pentatricopeptide repeat (PPR) and other helical repeat proteins, and RNA editing and splicing factors. We found that the Arabidopsis chloroplast terminome is complex, and in some cases surprising. For example, both known and new transcription start sites (TSS) were identified, sometimes internal or antisense to known transcripts, and a general imprecision of both processed 5’ and 3’ ends was observed. To highlight the comparative potential of Terminome-Seq, we examined the *pnp1-1* mutant which lacks the major 3’ processing enzyme polynucleotide phosphorylase (33, 34), revealing a largely reshaped terminome. Overall, our results showcase Terminome-Seq as a valuable addition to the organelle gene expression analysis toolkit.

## MATERIALS AND METHODS

### Plant Material

*Arabidopsis thaliana* Col-0 and *pnp1-1* seeds were germinated on MS medium with 16 hrs of light per day at 23°C. Three-week old leaf material was flash-frozen in liquid nitrogen, and total RNA was isolated using TRI Reagent according to the manufacturer’s instructions (www.sigmaaldrich.com).

### Terminome Library Synthesis and analysis

All libraries were produced from 1 µg of DNase I-treated RNA (www.neb.com), and for TAP-treated samples, tobacco acid phosphorylase (TAP; www.epibio.com) was used according to the manufacturer’s instructions with heat inactivation at the end of the incubation period. Library synthesis was carried out using the Illumina TruSeq Small RNA library preparation kit (www.illumina.com) intended to capture the RNA population containing a 5’ phosphate and 3’ hydroxyl group. Minor modifications were made to the protocol depending on whether a native 5’ or 3’ end was the target. Libraries intended for native 3’ end capture followed the protocol with initial 3’ adapter ligation using T4 RNA ligase 2, a deletion mutant that can only ligate a 3’ hydroxyl group to a 5’ adenylated RNA, consistent with the 3’ RNA adapter chemistry. After ligation, the RNA was fragmented using a Covaris sonicator (www.covaris.com), with a target size of 200 nt, followed by ethanol precipitation for concentration and 5’ adapter ligation with T4 RNA ligase. Libraries intended for native 5’ end capture required further adjustments. The order of adapter ligation was reversed: 5’ adapter ligation (with T4 RNA ligase) – sonication – ethanol precipitation – 3’ adapter ligation (with T4 RNA ligase 2). In this case, excess 5’ adapter remaining following the sonication and ethanol precipitation could ligate to added 3’ adapter, but not to any new 5’ ends created through sonication as the new 5’ ends would not be adenylated. This resulted in unwanted adapter dimers that were preferentially amplified during library amplification (PCR1) due to their small size (∼133 bp). Therefore, size selection was performed on the products from PCR1, retaining only products over 200 bp using Pippin Prep (www.sagescience.com). A second PCR amplification (PCR2) was executed on these products. Quality control was performed after Pippin size selection and before library submission for sequencing using an Agilent BioAnalyzer (www.agilent.com). Details about the procedure and the Pippin Prep are available at https://github.com/BenoitCastandet/Terminome_Seq. The final cDNA libraries were purified using magnetic AMPure beads (www.beckman.com) following the manufacturer’s protocol. Multiple steps in the above protocol, including fragmentation, ethanol precipitation, Pippin size selection (5’ libraries only), and AMPure purification of cDNA libraries, resulted in a bias towards the retention, and therefore sequencing, of fragments >67 nt.

Libraries were pooled and sequenced on a NextSeq500 Sequencer (www.illumina.com) using the v3 kit, with paired-end reads generating 40 bp long R1 reads and 35 bp long R2 reads for all libraries. R1 reads are only of use for libraries generated to obtain 5’ related data, while R2 reads contain data related to 3’ ends and therefore are only relevant for libraries generated to obtain 3’ data. The detailed pipeline used to analyze the relevant reads is available at https://github.com/BenoitCastandet/Terminome_Seq. Briefly, the quality of relevant reads was checked using fastq-mcf (https://github.com/ExpressionAnalysis/ea-utils/blob/wiki/FastqMcf.md) followed by alignment to the chloroplast reference genome, Arabidopsis TAIR10 version modified to add the first exon of the chloroplast gene *ycf3*, using tophat2 (https://ccb.jhu.edu/software/tophat/index.shtml). Two customized scripts allowed us to extract the positions of the 5’ and 3’ termini and the results were normalized according to the numbers of reads aligned to the chloroplast genome. Normalized data are available in Supplementary Table S1.

### 5’ RACE

Total RNA was isolated from mature leaf tissue using TRI Reagent® and treated with DNase I (Ambion; http://www.thermofisher.com). 5’ Rapid Amplification of cDNA Ends (RACE) reactions were completed as described (35) with some modifications. For analysis of primary transcripts, 4 µg of RNA was treated with TAP (0.5U/4 µg RNA; Epicenter, http://www.epibio.com) for 1 hr at 37°C with RNaseOUT (40U; Invitrogen, http://www.thermofisher.com), followed by phenol/chloroform extraction and ethanol precipitation. The 5’ RACE adapter (Supplementary Table S2) was then ligated to the 5’ ends of the + or –TAP treated RNA with T4 RNA ligase (Ambion). For this, 10 µM of 5’ RACE adapter and 4 µg of +/–TAP-treated RNA were incubated for 5 min at 65°C. Samples were chilled on ice and then 5U of T4 RNA ligase, 1x ligase buffer, 1 mM ATP and 40U of RNaseOUT were added, and the reactions were incubated for 1 hr at 37°C. Ligated RNA was phenol/chloroform purified and precipitated with ethanol, and cDNA was generated using SuperScript III (Invitrogen) with random hexamers according to the manufacturer’s protocol. Transcripts were amplified by PCR using the 3’ gene-specific primers indicated in the Figure Legends and the 5’ RACE primer (Supplementary Table S2). Amplicons were visualized after separation through a 1% agarose gel and gel purified. Purified bands were used for a second, nested PCR reaction with the indicated 3’ RACE primer and the RACE 5’ nested primer to ensure amplified sequences were specific to the intended target and/or increase amplicon intensity as the bands were directly sequenced, or cloned and sequenced. The nested PCR reactions were visualized in 1% agarose gels, and specific bands were gel purified, cloned and sequenced.

## RESULTS

### Terminome Sequencing Strategy and Overview

Identification of RNA termini has traditionally been performed by sequencing individual clones from RACE products, primer extension, or nuclease protection assays. A recent improvement was the substitution of high-throughput sequencing for product-by-product analysis (36–40). Because of their focus on nucleus-encoded RNAs, these protocols rely on the presence of 3’ poly(A) tails, rendering them unsuitable for studying organellar RNAs where such tails mark transcripts for degradation (27, 41, 42). Our experimental design included the sequencing of three distinct libraries devoted to the identification of both primary and processed 5’ ends, or 3’ ends. The protocol (Supplementary Figure S1) was modified from the Illumina TruSeq Small RNA kit, which relies on sequential ligation of adaptors prior to cDNA synthesis and amplification. For Terminome-Seq, the initial ligation of 5’ or 3’ adapter captures the native 5’ and 3’ termini, respectively, then the RNA is fragmented followed by a second adapter ligation step before amplification. To differentiate and identify primary transcripts that correspond to transcription start sites (TSS), RNA was pretreated with TAP prior to 5’ adaptor ligation. TAP converts triphosphorylated 5’ ends, uniquely found in primary transcripts, to monophosphates, making them amenable to adapter ligation. Thus, libraries created with or without TAP pretreatment (+TAP and –TAP) can be compared to identify TSS. Duplicate libraries were constructed from *Arabidopsis thaliana* Col-0 biological replicates and the results show high correlation (Supplementary Figure S2). Native transcript ends were defined as the first nucleotide of each read, following suitable manipulation of the primary sequencing. The full coverage for Terminome-Seq at single nucleotide resolution for 5’ (+/–TAP) and 3’ ends generated for this manuscript can be accessed in Supplementary Table S1. It is important to note that transcripts with post-transcriptional modifications, such as tRNAs with a post-transcriptionally added CCA tail, or RNA degradation intermediates bearing polynucleotide tails, would not fully align to the genome and are therefore excluded from our curated dataset.

A genome-wide view of end distribution is depicted in Figure 1. Clusters of 5’ and 3’ ends, covering a remarkable 13% and 16% of the genome, respectively, are clearly discernable, but are largely absent from areas antisense to known transcripts. In contrast, the full chloroplast genome is covered by reads from total RNA-Seq experiments (2, 19, 20, 23). However, when considering only positions representing at least 0.001% of total reads (10 reads per million, or RPM), less than 0.7% of the genome (1,628 processed 5’ ends and 1,299 3’ ends, respectively) was represented (Figure 2A). Thus, the vast majority of termini are stochastically found at low levels, probably representing RNA degradation products or processing intermediates.

**Figure 1:**
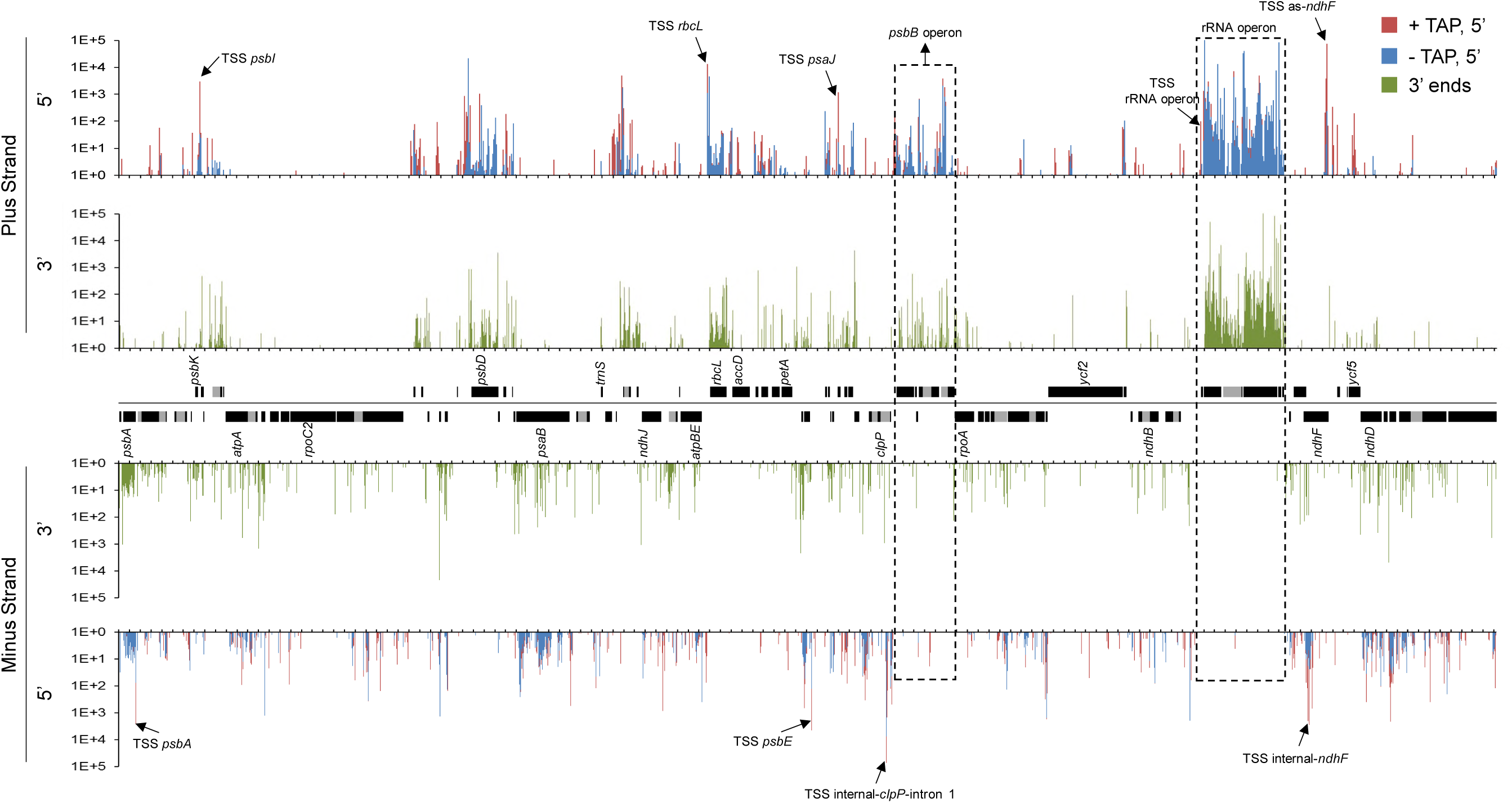
Plastome-scale view of Terminome-Seq results. End coverages are the average of two Col-0 biological replicates and given in RPM. 5’ ends obtained with or without TAP treatment are red and blue, respectively, and 3’ ends are displayed in green. Gene models are indicated between the tracks corresponding to the plus and minus strands of the plastome. One copy of the large inverted repeat is omitted for clarity. Selected TSS described in more detail in the main text are marked by arrows and labeled, as are the *psbB* and rRNA operons. Tick marks are every 1,000 nt.

**Figure 2:**
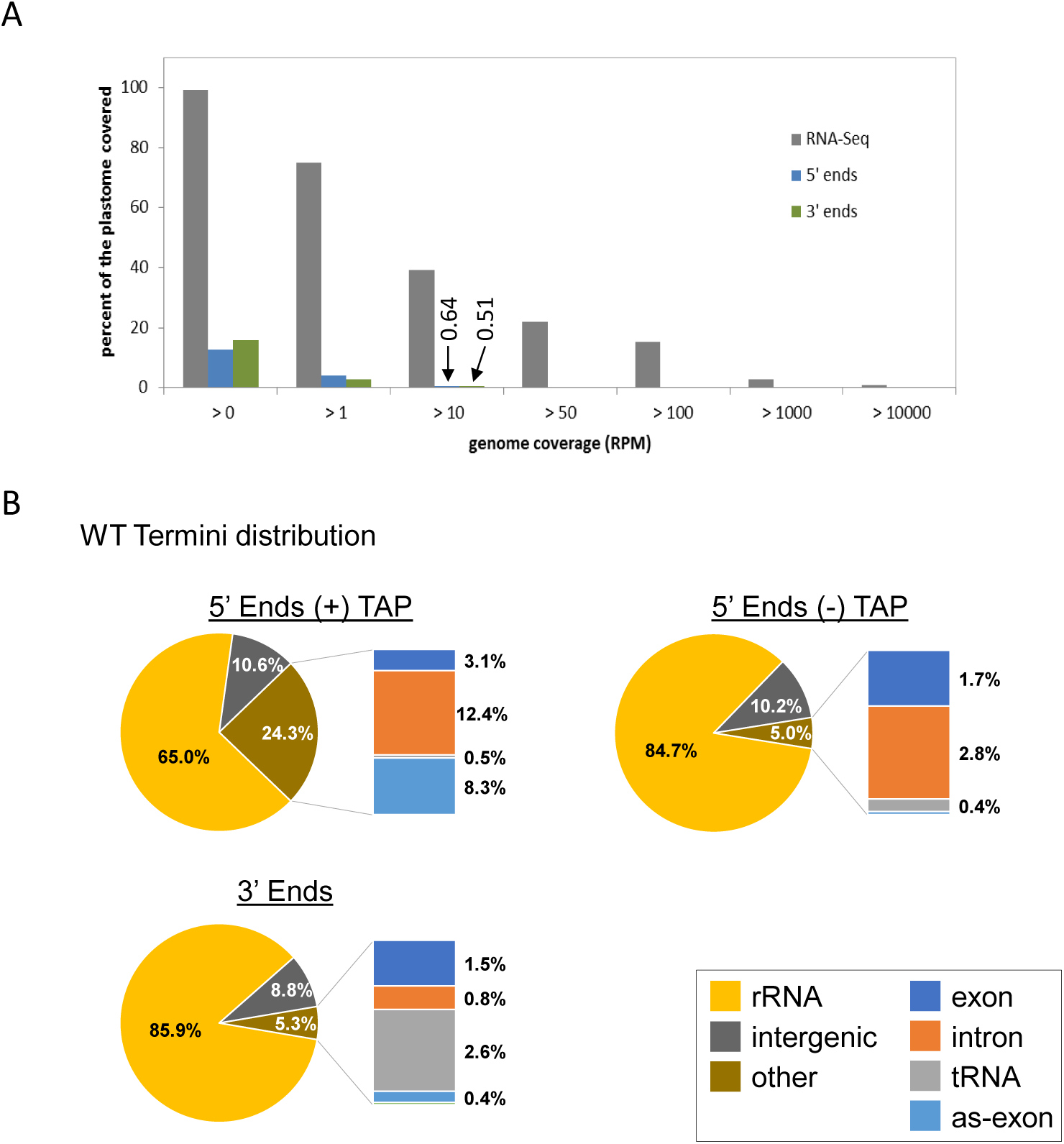
Coverage and distribution of transcript termini. **A)** Comparison of genome coverage between RNA-Seq and Terminome-Seq. While the plastome is almost fully covered by at least one read in RNA-Seq, only 12.7% and 15.8% is covered by 5’ and 3’ ends, respectively. Data for RNA-Seq correspond to the average of two previously published WT replicates (19). Termini coverage at >10 RPM is marked and discussed in the text. **B)** Terminome-Seq read distribution in the WT. The results are the average of two biological replicates. Reads antisense to exons (as-exon) refers to reads mapping to the antisense strand of a known coding region.

To compare the abundance of termini across regions of the chloroplast genome, we informatically divided them into six types: rRNAs, tRNAs, exons (corresponding to protein-coding regions), introns, intergenic regions (anything else), and positions on the antisense strand of exons. When categorized this way, Also notable in Figure 2B shows that approximately 85% of the 5’ and 3’ ends were found in the rRNAs. The high degree of rRNA transcript ends was expected due to the elevated expression level of the rRNA operon and lack of rRNA depletion during the library preparation. Addition of an rRNA depletion step could help to map low abundance transcript ends that may have been overlooked in this study. Ends corresponding to intergenic regions were overrepresented compared to exons (Figure 2B). Indeed, even though intergenic regions cover around 25% of the genome (43), they contained 10.2% and 8.8% of the –TAP 5’ and 3’ ends, respectively, whereas only 1.7% and 1.5% of the ends mapped within exons. TAP treatment, which allowed TSS to be represented, significantly altered these proportions, with the proportion of 5’ ends mapping to rRNAs decreasing to 65%, mainly in favor of an increase in reads mapping to introns (12.4%) and antisense to exons (8.3%; Figure 2B). Remarkably, an overabundance of TSS is found in the first *clpP* intron (TSS internal-*clpP*-intron 1) and antisense to the *ndhF* transcript (as-*ndhF*; highlighted in Figure 1).

### The chloroplast genome contains at least 215 TSS

Because the 5’ ends of most primary transcripts are marked by a 5’ triphosphate, they should be highly overrepresented in +TAP-treated versus –TAP libraries. We therefore defined a TSS as any 5’ end that was at least 10 times more abundant in +TAP libraries, a calculation that generated 352 positions (Figure 3A). Because of the generally precise mechanism of transcription initiation, a true TSS should be a single predominant nucleotide rather than a cluster. We therefore filtered out putative TSS where the first nucleotide (the TSS itself) represented less than 50% of the coverage over a 5-nt stretch where it was the first nucleotide, as was done for barley (20). This reduced the number of TSS being considered to 215, with the others potentially representing stochastic initiation (Supplementary Table S3). Among the 215 defined TSS, 81% of the initiating nucleotides were purines (119 A’s and 55 G’s; Figure 3A). This trend is consistent with barley, where purines defined >80% of TSS (20). Our data identified 45 previously-described Arabidopsis TSS, 16 of which are in orthologous positions in barley (20), meeting our expectations considering that more closely-related species like *Arabidopsis thaliana* and *Nicotiana tabacum* differ in their promoter usage for some genes (45).

**Figure 3:**
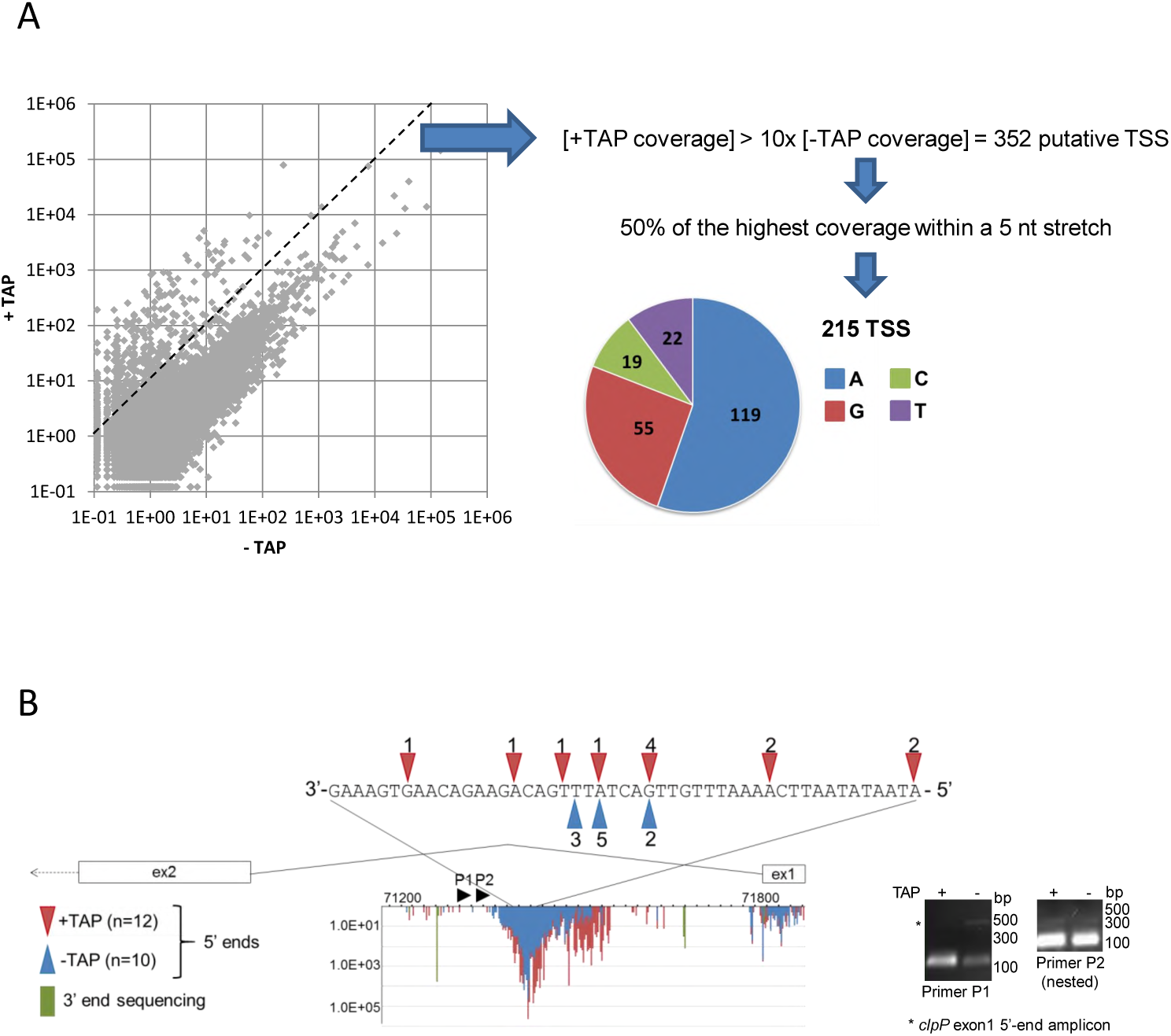
TSS analysis. **A)** The abundance of 5’ ends at each position for both genome strands was compared between +TAP and –TAP libraries. The dashed line separates the 352 ends that have a +TAP/–TAP ratio >10 from those with a lower ratio. Putative TSS were filtered to remove any ends that did not reach 50% of the coverage of the most represented read within a 5 nt stretch. The pie chart graphs the initiating nucleotide of the remaining 215 TSS. **B)** The novel TSS detected within *clpP* intron 1 were mapped using 5’ RACE. 5’ RACE was completed with (red) and without (blue) prior TAP treatment, and sequenced clones are represented by colored arrowheads above/below the nucleotide sequence. The stained gel of the corresponding PCR reactions is shown at right. The gene model between exons 1 and 2 is shown with Terminome-Seq results. +TAP 5’ ends are in red, –TAP ends in blue, and 3’ ends are in green. The X-axis is genomic position and the Y-axis is RPM coverage. Black arrowheads P1 and P2 represent the 3’ primers used for 5’ RACE.

Most plastid genes belong to polycistronic units and have classically been described as being co-transcribed. A few genes, such as *psbA, rbcL* and *ndhF*, are considered to be monocistronic, which is also the tendency for tRNAs outside the rRNA operon (46, 47). TSS could be identified at the 5’ ends of all but one of the 20 main transcriptional units, with *petL-petG-psaJ-rpl33-rps18* being the exception. The processed *petL* 5’ end is easily identifiable, however (see below). This suggests rapid maturation of the primary transcript, a phenomenon that is more prevalent in *Chlamydomonas reinhardtii* chloroplasts, where a recent transcriptome analysis revealed only 23 TSS, albeit using an entirely different method (48). Terminome-Seq also confirmed our earlier discovery of TSS upstream of *trnC, trnF* and *trnN* (22). All transcriptional units also contained internal TSS, similar to barley (20), which could reflect mechanisms to allow for differential expression of clustered genes within a larger cluster as is particularly well-documented for the *psbD-C* cluster segment of the barley *psbK* gene cluster (49).

TAP treatment revealed three genomic areas with unexpectedly massive numbers of initiation events. As mentioned above, these lie in the first *clpP* intron, and on the sense and antisense strands of *ndhF* (Figure 1 and Supplementary Table S3). TSS antisense to *ndhF* are distributed over nearly 300 nt, and some of them were also characterized in barley and tobacco (20, 50). The TSS internal to *clpP* intron 1 and on the sense strand of *ndhF* are distributed over approximately 160 and 600 nt, respectively, and to our knowledge have not been previously described. As a validation step, we confirmed that at least some of these ends could be amplified by 5’ RACE following TAP treatment (Figure 3B and Supplementary Figure S3A).

Although we attempted to obtain an exhaustive catalog of TSS, the data filters we used to reduce the false discovery rate eliminated a few known initiation sites (Supplementary Figure S3B). For instance, the P*psbN* -32 promoter (51) and P*atpE* -430 (52, 53) are absent from our list because they have a +TAP:–TAP ratio of <10 (9.9 for P*atpE* and 2.5 for P*psbN*). On the other hand, our data shed light on the uncertainty of whether P*psbD* -186 (position 32525) is a genuine promoter or processed end (54–56), with our results being strongly diagnostic of a post-transcriptional processing site. The opposite is true for the *ndhA* -66 (position 122076) 5’ end that has been described as being created post-transcriptionally in maize through the action of SOT1 PPR53 (57), but is a TSS based on our data.

### The case of the *psbB* operon

The *psbB* operon is particularly well studied, containing five genes on the plus strand (*psbB-psbT-psbH-petB-petD*, the last two containing introns) and one on the minus strand (*psbN*). Altogether, over 20 accumulating transcripts generated from this operon have been characterized (5, 10, 58), making it an attractive testbed for the ability of Terminome-Seq to reflect such a complex landscape accurately.

Figure 4A shows the positions of the eight TSS (numbered in red), along with at least 12 processed 5’ ends (numbered in blue) and 17 3’ ends (numbered in green), which are annotated in Table 1. A consistent feature of 5’ ends is their clustered organization around a dominant peak, whereas the 3’ termini are more discrete. Heterodisperse 5’ ends are reminiscent of degradation intermediates, perhaps created by an enzyme such as RNase J that would progressively stall as it encountered secondary structures or bound proteins. 3’ ends that were more discrete could be identified in both coding and intergenic regions, probably representing a mix of degradation intermediates and mature 3’ ends. Among the processed 5’ ends and 3’ termini found by Terminome-Seq, several had been previously described. These include the 5’ ends *psbB* -51 (58), *psbH* -44 (59, 60) and -67 (61), and *petB* -47 (62, 63), as well as the 3’ termini *psbT* +60 (30), +223 (61), *psbH* +109 (62, 63), *petB* +67 (64), *petD* +94 (65), and *psbN* +39 (30). Thus, there was excellent overlap between our data and previously published work.

**Table 1.**
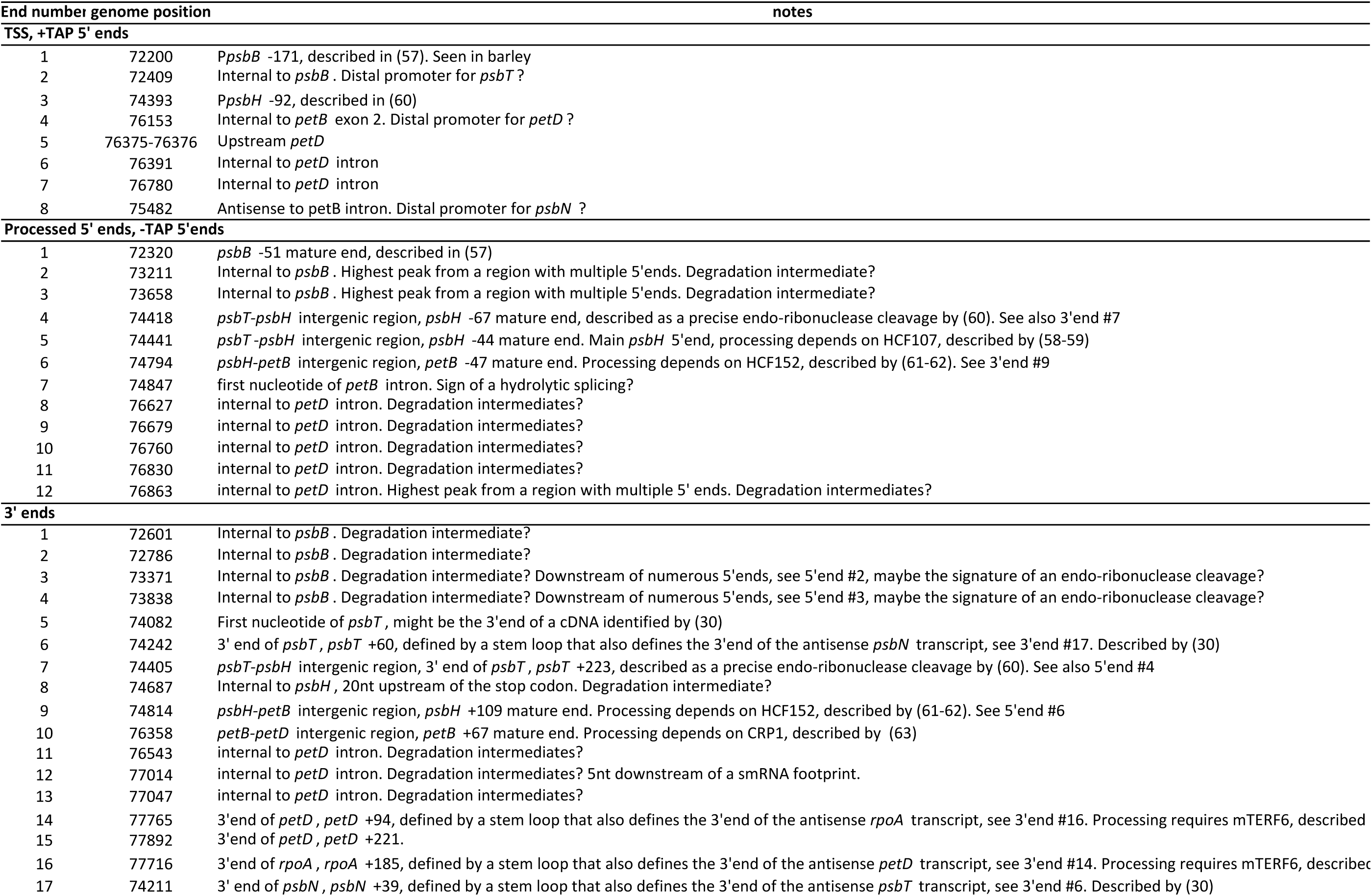
Decsription of transcript ends originating from the *psbB* operon.

**Figure 4:**
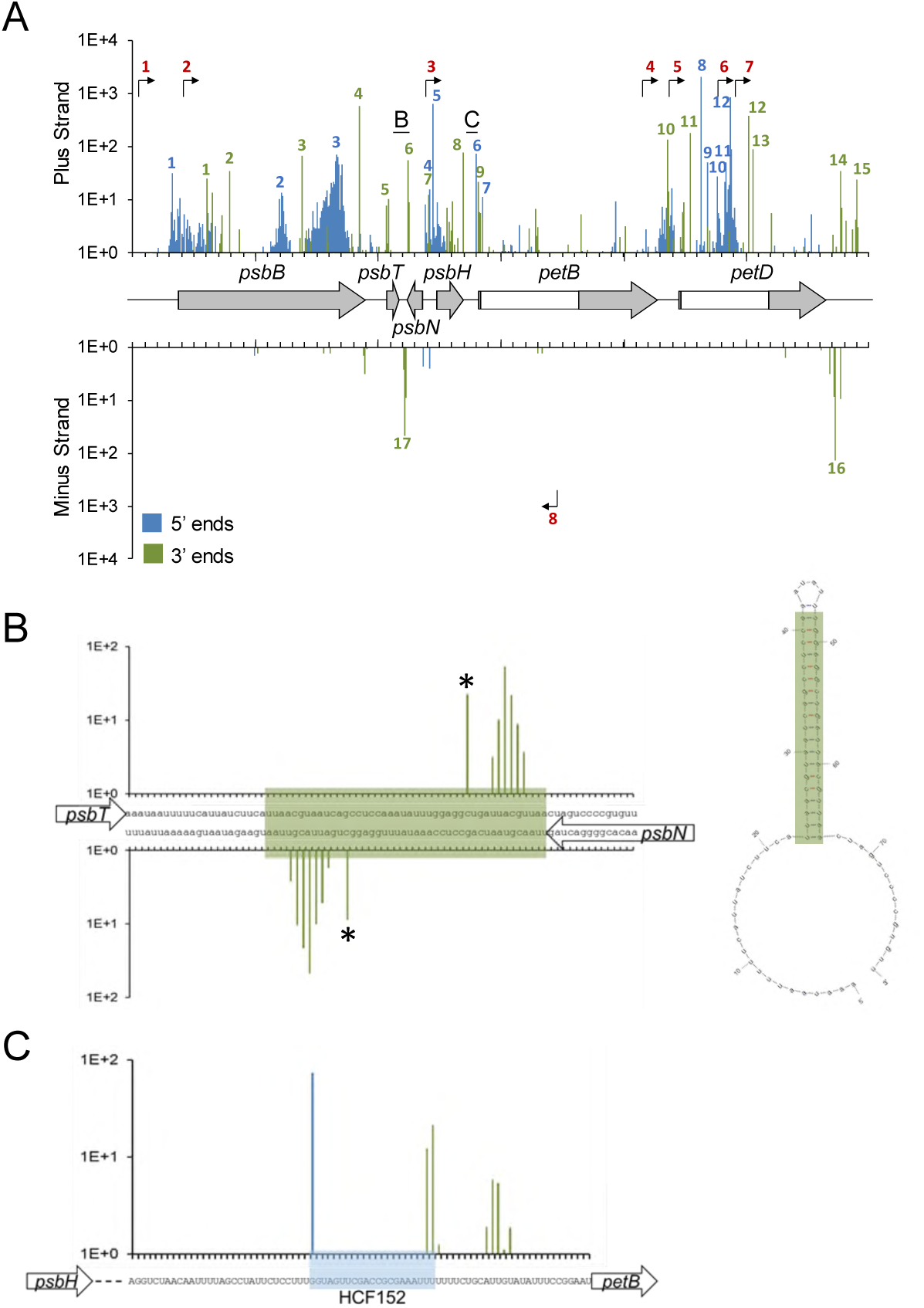
Transcript termini of the *psbB* operon highlighting the role of a secondary structure and RNA binding protein in defining ends. **A)** Terminome-Seq coverage of the *psbB* operon with the corresponding gene models, with exons in gray and introns in white. –TAP 5’ ends are in blue and 3’ ends are in green; bent arrows represent TSS inferred from +TAP data. Underlined letters mark the locations that are expanded in panels B and C; numbered peaks and promoters refer to features listed in Table 1. **B)** 3’ end coverage for a stem-loop structure between *psbT* and *psbN*. The stem is highlighted in green in the nucleotide sequence and the Mfold (123) predicted secondary structure is at right. Asterisks highlight the previously described ends (30). **C)** The gene model, nucleotide sequence, and end coverage for the HCF152 binding site. Reads accumulate at both the 5’ and 3’ ends of the binding site on the plus strand, indicative of a protected RNA fragment. The color code is the same as in panel A.

We conducted a similar analysis for the *atpI-atpH-atpF-atpA* and *ndhH-ndhA-ndhI-ndhG-ndhE-psaC-ndhD* gene clusters (Supplementary Figure S4). Five TSS were identified in the *atpI* cluster, including what might be specific promoters for *atpH* and *atpA*, and 15 TSS in the *ndhH* cluster, which were located predominantly toward the distal end. Most of these 3’ and 5’ termini were previously identified in a thorough investigation of the *atpI* cluster processing (53). The existence of an accumulating monocistronic *atpI* transcript in Arabidopsis has been debated (53, 66) and our data suggest that the main *atpI* 3’ end is 584 nt downstream of the stop codon, inside *atpH*. This 3’ end is more abundant than the one at position +493, whose processing depends on PPR10 (53, 67). 27 termini in addition to TSS were found for the *ndhH* cluster. No detailed information was previously published on the transcript population, however it is quite complex based on gel blot analysis (68), in keeping with our results.

### The roles of RNA secondary structures and RNA-binding proteins in shaping the terminome

Chloroplast RNA termini are known to be stabilized by stem-loop structures, as well as by sequence-specific and general RBPs (6, 7, 69). Both of these mechanisms act on transcripts from the *psbB* operon, for which termini were described above. While the strategy used here to make the terminome libraries would be biased against smRNAs, the processing sites from longer, precursor RNAs could be detected. The *psbT-psbN* intergenic region, for example, forms a stem-loop that defines the 3’ ends of transcripts encoded on opposite strands (30). Terminome-Seq confirmed the previously identified 3’ ends inside the stem and additionally showed staggered 3’ ends closer to the base of the stem, an expected phenomenon given the tendency of exoribonucleases to stall at such positions (Figure 4B). Another stem-loop, originally described as a “twin terminator” in spinach (70), defines the 3’ ends of the *petD* and *rpoA* transcripts encoded on opposite strands, and a similar structure exists for *petA* and *psbJ* (Supplementary Figure S5). The co-existence of two sets of 3’ ends associated with the *psbT* - *psbN* stem-loop is also true for the transcripts of *rbcL* (ends in positions 56485 and 56488; see below) and *psbA* (ends in positions 293 and 285; Supplementary Figure S5), probably reflecting breathing in AU-rich regions of the secondary structures.

The *psbH-petB* intergenic region is known to be bound by HCF152 (62, 63), a PPR protein that defines the *psbH* 3’ and *petB* 5’ ends (71). In agreement with these results, Terminome-Seq data analysis identified a single major 5’ end for *petB* correlating with the 5’ end of a smRNA established as the footprint protected by HCF152 binding (9, 67, 72). The Terminome-Seq *psbH* 3’ end also correlates with HCF152 binding, with an additional cluster of 3’ ends found ∼10 nt downstream, possibly reflecting the different stalling characteristics of the exoribonucleases PNPase and RNase II (Figure 4C).

The correlation between RBPs, their footprints found in smRNAs, and Terminome-Seq ends can be extended beyond HCF152 to at least 13 RBPs that have been described as involved in RNA maturation (Supplementary Figure S6). These include PPR10, which protects the two adjacent transcripts *atpI* and *atpH* and assists in the processing of both their 5’ and 3’ ends (67, 71, 73); and HCF107 and CRP1 which, like HCF152, target the *psbB* operon, in particular the *psbH* -44 5’ end and the *petB* +67 3’ end. We can further generalize this phenomenon, since, on a plastome-wide level, termini are enriched in areas containing smRNAs, which are in many cases likely to be marks of RBPs that still await identification (Supplementary Table S5).

### PNPase deficiency has broad impacts on both 5’ and 3’ termini

The roles of PNPase in chloroplast transcript 3’ maturation, intron degradation, and tRNA processing, as well as an ancillary role in phosphorus metabolism, have been well documented (22, 33, 34, 74–76). Although RNA-Seq shows that most chloroplast RNAs contain 3’ extensions in the *pnp1-1* null mutant (22), the individual termini were not systematically compared between WT and mutant. Such a comparison could reveal more precisely the types of sequences and structures that impede PNPase activity, and highlight its overall impact on the plastid terminome. We proceeded to analyze duplicate *pnp1-1* Terminome-Seq libraries, and decided against performing TAP treatment to capture primary 5’ ends because PNPase is not known or expected to affect transcription initiation.

A plastome-wide view of *pnp1-1* termini is presented in Figure 5A. While the WT and *pnp1-1* patterns show substantial overlap, there are numerous areas where reads are specific to *pnp1-1*. This is quantified as genome coverage of reads (Figure 5B) where, regardless of the RPM threshold applied, coverage is higher in *pnp1-1* than in WT. In aggregate, 5’ end coverage increases from 12.7% to 26.1%, and 3’ coverage from 26.1% to 39.5%. At When a threshold of >10 RPM is applied to remove low abundance termini, coverage is approximately 1% in the mutant, with 2,420 5’ termini and 2,744 3’ termini exceeding the threshold. This total of 5,164 >10 RPM termini in *pnp1-1* represents an increase of 76% over the combined vs. 2,927 termini found in the WT. The locations of termini are also dramatically altered between genotypes (compare Figures 2B and 5C). Another noticeable difference is the decline in the proportion of termini found in rRNA, which can be rationalized as a relative increase in non-rRNA termini in the mutant. In the –TAP 5’ termini population, the mutant differs by having an increase in intronic ends and a decrease in tRNA ends, the latter of which holds true for 3’ termini as well.

**Figure 5:**
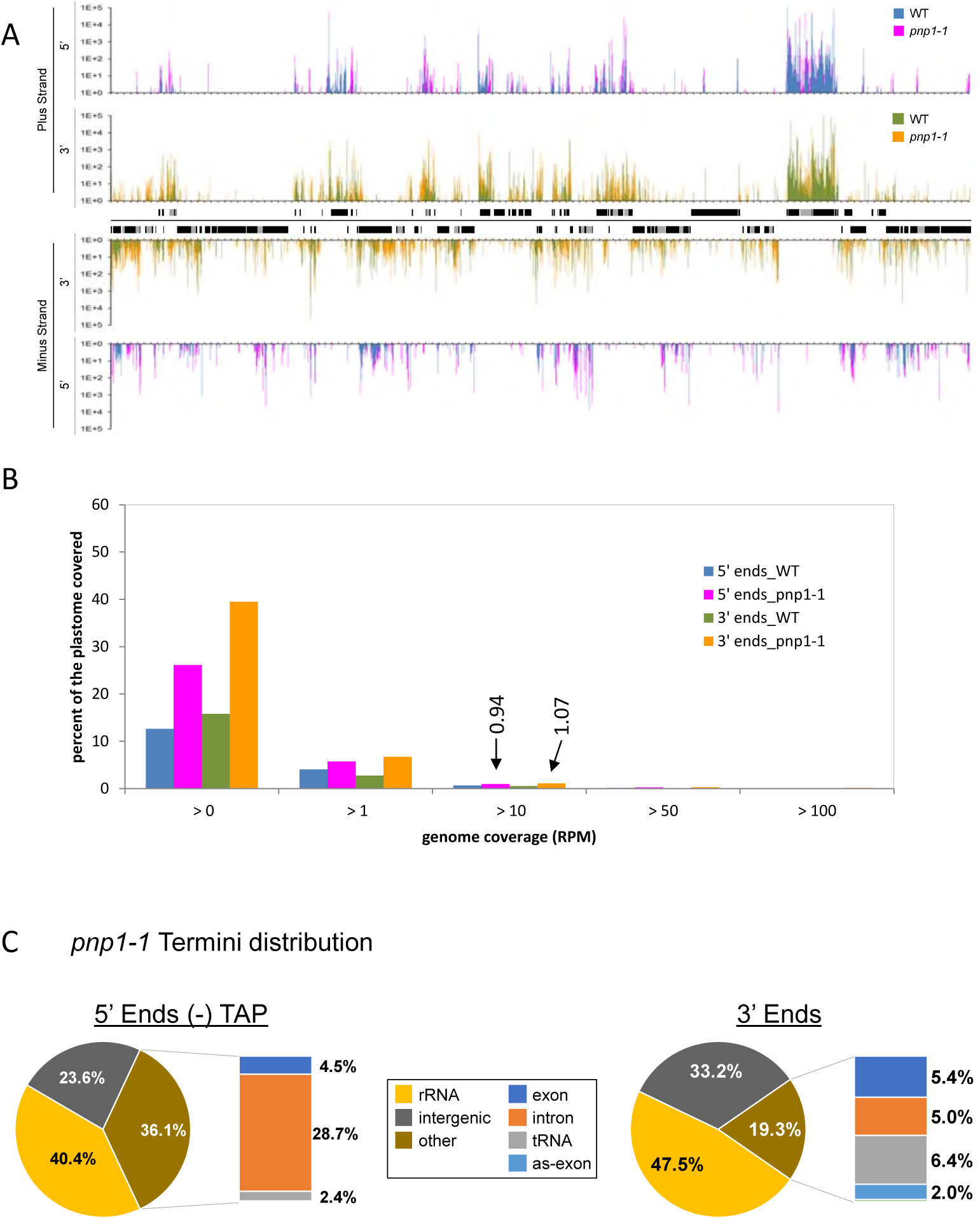
Distribution, coverage and location of transcript termini in *pnp1-1*. **A)** Plastome-scale view of end coverages from the average of two Col-0 and *pnp1-1* biological replicates in RPM. 5’ ends obtained without TAP treatment are blue (WT) and pink (*pnp1-1*), and 3’ ends are displayed in green (WT) and orange (*pnp1-1*). Gene models are indicated between the tracks corresponding to the plus and minus strands of the plastome. One copy of the large inverted repeat is omitted for clarity. Tick marks are every 1,000 nt. **B)** Comparison of Terminome-Seq coverage for WT and *pnp1-1*. While 12.7% and 15.8% of the WT plastome is represented by 5’ and 3’ ends, respectively, these numbers increase to 26.1% and 39.5%, respectively, in *pnp1-1*. Termini coverage at >10 RPM (0.94% and 1.07% for 5’ and 3’ ends, respectively, in *pnp1-1*) is marked and discussed in the text. **C)** Terminome-Seq read distribution in *pnp1-1*. The results are the average of two biological replicates. as-exon refers to reads mapping to the antisense strand of known coding regions.

A more pronounced effect on 3’ vs. 5’ ends in *pnp1-1* can be gleaned from Figure 6A, which shows that 349 5’ termini and 1,348 3’ termini are at least 10 times more abundant in the mutant compared to the WT, whereas only 96 and 150, respectively, decrease at least ten-fold in the mutant. The tendency of termini to accumulate in the mutant is in keeping with the degradative function of PNPase, with the expected bias toward 3’ termini. The strong effect of PNPase on the 5’ terminome was unexpected and somewhat counterintuitive, given that it is a 3’ to 5’ exonuclease. We have previously shown, however, that PNPase degrades tRNA 5’ leader sequences following their liberation by RNase P cleavage (22), and Terminome-Seq based evidence of this phenomenon is shown in Supplementary Figure S7. We speculate that other 5’ termini that hyperaccumulate in *pnp1-1* represent the upstream termini of other endonucleolytic cleavage products that are usually processed removed by the 3’→5’ activity of PNPase.

**Figure 6:**
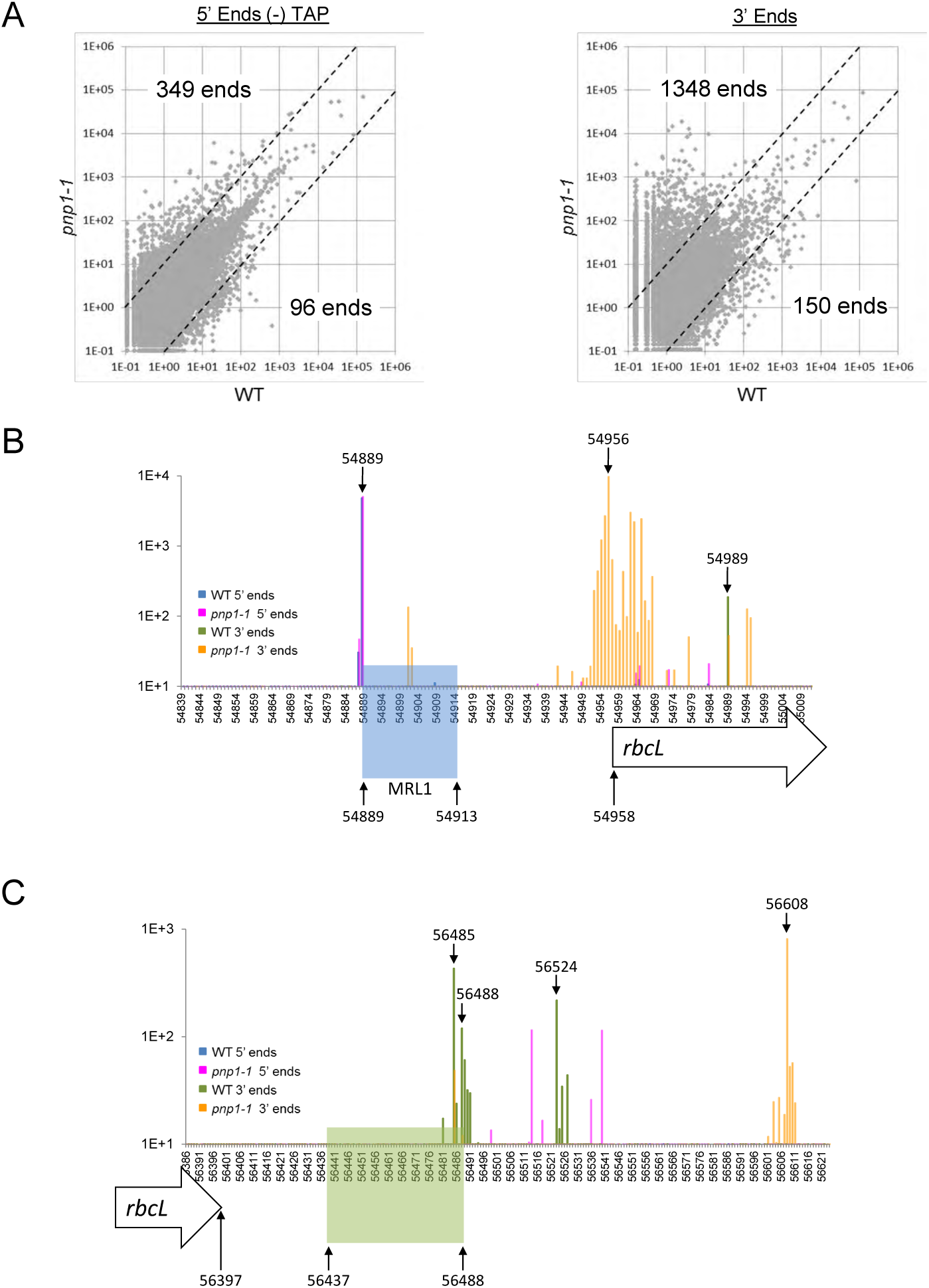
Terminome-Seq coverage in *pnp1-1*. **A)** The RPM abundance of –TAP 5’ ends and 3’ ends was compared between WT and *pnp1-1* at a genomic level. The dashed lines separate ends that are at least 10-fold more abundant in a given genotype. For example, 349 5’ ends are more abundant in the PNPase mutant. **B)** Terminome-Seq coverage upstream of the *rbcL* gene. Color coding of ends is provided in an inset. Genome position 54958 is the *rbcL* coding region 5’ end according to the TAIR10 annotation. The *rbcL* processed 5’ end (position 54889) correlates with the 5’ end of the smRNA footprint of MRL1 (highlighted in blue). **C)** Terminome-Seq coverage downstream of the *rbcL* gene, with labeling as in Panel B. Genome position 56397 is the 3’ end of the coding region. The stem-loop downstream of the gene (positions 56437-56488) is highlighted in green and matches a smRNA (67). Other genome positions discussed in the text are also labeled.

To illustrate the utility of Terminome-Seq for characterizing a ribonuclease mutant at the individual gene level, results are shown for the monocistronic *rbcL* transcript (Figures 6B and 6C), which in plants accumulates as two species with primary and processed 5’ ends. The processed 5’ end is protected by the PPR protein MRL1, which prevents degradation by RNase J (77, 78). MRL1 leaves a smRNA footprint (Figure 6B, blue shading), whose 5’ end is represented as similarly abundant termini in WT and *pnp1-1* at genome position 54889. There is, however, a cluster of 3’ ends located ∼40 nt downstream of the MRL1 binding site (with a peak at position 54956) that is present only in the mutant, even though there is a less abundant 3’ end slightly downstream (position 54989) in both genetic backgrounds. This indicates that PNPase likely degrades *rbcL* mRNA until it is stalled by MRL1 and/or nearby RNA secondary structures, as such termini are absent in the WT. The 3’ end at position 54901 may be an RNase II stall site, given the known cooperation of RNase II and PNPase in 3’→5’ RNA decay (79). A comparison of WT and *pnp1-1* ends in the vicinity of all described RBP sites is provided in Supplementary Figure S6.

At its 3’ end, the *rbcL* transcript is defined by a highly conserved stem-loop (80, 81) represented as a smRNA (67, 72). RNA-Seq showed higher coverage in *pnp1-1* compared to the WT beginning about 10 nt downstream of this structure (22). Using Terminome-Seq results, we identified three different 3’ ends in WT plants, two of them directly at the 3’ base of the stem (positions 56485 and 56488) and one 36 nt downstream (position 56524; Figure 6C). The 3’ end at position 56485 could also be identified in *pnp1-1*, although it was less abundant than in WT, suggesting that its production is not fully dependent on PNPase. On the contrary, ends at positions 56488 and 56524 are certainly produced through the action of PNPase because they are missing in the mutant. The most abundant 3’ ends in the mutant cluster around position 56608, 120 nt downstream of the stem-loop (Figure 6C), accounting for the 3’ extension previously noted in RNA gel blots, which represents the stall point of RNase II (79). Terminome-Seq evidence for several other putative 3’ extensions is presented in Supplementary Figure S8.

Finally, we present an overview of rRNA operon ends for the WT and *pnp1-1* (Supplementary Figure S9). This analysis revealed termini corresponding to known processing sites (6) for all four rRNAs, and also showed clear peaks in *pnp1-1* for the well-known 23S rRNA extension in this mutant (82). Of note, the first hidden break of the 23S rRNA was not clearly delineated compared to the second 23S rRNA hidden break, in keeping with previously mapped ends in this region (19). Surprisingly, in the 5’ part of 16S rRNA there were numerous, staggered 5’ termini indicative of a processing or degradation event, albeit at 1-2 logs lower abundance than the mature 5’ and 3’ ends. RNA processing at this position, however, has not been previously described and further analysis of this region is needed to evaluate the origins of these termini.

## DISCUSSION

### The chloroplast TSS landscape

In this work, we systematically sequenced *Arabidopsis thaliana* chloroplast RNA 3’ termini as well as primary and processed 5’ ends, using TAP treatment to discriminate between them. Such strategies are under continual improvement, as transcriptome analyses generally endeavor to be as comprehensive and quantitative as possible. In our case, transcript chemistry, size and secondary structure will all introduce biases to the dataset ultimately used to draw the main conclusions. In addition, factors such as ligation bias – the preferential ligation of adapters to certain sequences (83, 84), undoubtedly impacted the selection and ratios of the transcript ends we have reported. Therefore, our conclusions – particularly quantitative ones – should be taken with those caveats in mind.

The strategy as implemented led to the description of 215 TSS meeting defined expression thresholds, which are widely distributed in the plastome (Figure 1) and is consistent in number with the 176 TSS mapped in mature barley leaves (20). These TSS are created by two RNA polymerase types, a bacterial-like, plastid-encoded RNA polymerase (PEP) and two phage-like, nucleus-encoded, RNA polymerases (NEP). NEP and PEP operate simultaneously, however our samples were taken from tissue populated by mature chloroplasts, where PEP activity predominates (85). Since NEP transcription primarily occurs during early differentiation, the TSS we describe likely underestimate the number of promoters utilized over the course of chloroplast differentiation. In barley, the use of the *albostrians* mutant with sectors lacking PEP allowed the discovery of 254 additional NEP-dependent TSS, giving a total of 398 unique TSS when overlap between PEP and NEP TSS was considered. At the same time, NEP is known to become promiscuous when PEP has been eliminated genetically (1, 86). The proportion of NEP promoters active under these circumstances that are also utilized in WT plants remains to be determined.

If we assume a total of ∼400 TSS in Arabidopsis when taking PEP and “developmentally hidden” NEP promoters into account, this would average to a TSS every ∼600 nt when both strands, but only one of the large inverted repeats, are used as a basis. This frequency is not surprising, and such phenomenon occurs in bacteria as well (87, 88). Given the AT-rich content of the plastid genome, functional PEP promoter -10 elements are likely to be present by chance, along with the short and highly variable elements that seem to constitute NEP promoters. The average 600 nt spacing we calculate would be reduced if we changed the 10-fold ratio threshold used to define TAP vs. non-TAP-dependent transcripts. Some known TSS such as P*atpE* -430, P*psbN* -32 and P*psbD* -948 (89), have a ±TAP ratio <10 and were therefore not formally considered to be TSS (Supplementary Figure S3B). Perhaps the biggest surprise from this Terminome-Seq dataset is the evidence for massive transcription initiation activity in the 3’ part of the *ndhF* gene, antisense to *ndhF*, and within the first intron of *clpP*. The former two areas represent a staggering ∼20% of the reads derived from primary transcripts, and could have functional implications in the chloroplast.

### Do some TSS mark transcripts with novel coding potential?

Chloroplasts are known to contain plastid non-coding RNAs (pncRNAs), with >100 in Arabidopsis (90), with functions remaining mostly unclear. Although many pncRNAs appear to be generated by read-through from adjacent genes, some are known to be primary transcripts (20, 23), and the high number of TSS identified here potentially increases the count of pncRNA promoters. For example, the internal *ndhF* TSS (Supplementary Figure S3A) probably initiates a 300-600 nt RNA that ends at the *ndhF* termini 110331 and 109924, the latter position being protected by an RBP or stem-loop since a smRNA from this position was identified (67). This pncRNA overlaps *ycf1* on the antisense strand and was previously thought to originate from *ndhF* readthrough (23). Initiation antisense to *ndhF* was also described in tobacco (91) and more recently in barley (20), and is likely responsible for additional Arabidopsis pncRNAs (nc89 and nc90) which were validated by RT-PCR and gel blots (23). The tobacco *ndhF* antisense TSS was proposed to be a proximal TSS for the downstream *rpl32* gene (91), which might also be the case in Arabidopsis.

Many pncRNAs are antisense to coding sequences, and for a few there is evidence they exert a regulatory function at the RNA level (19, 29, 61). On the other hand, many pncRNAs contain small open reading frames (ORFs) and could therefore encode unknown chloroplast proteins. Such ORFs were discarded from the chloroplast genome sequences first obtained from tobacco and *Marchantia* (92, 93), specifically ORFs shorter than 70 nt unless the products were known. In contrast, the functions of the longer, conserved, hypothetical coding frames (*ycfs*) were still discussed. Whether pncRNA-encoded ORFs are represented in the proteome remains to be determined, however data from bacterial and animal systems suggests that at least some antisense or intergenic RNAs harbor hidden genetic functions (94–96). In this case, the retention of TSS would be expected.

Among the better-studied YCFs, our data shows that *ycf15* (also annotated as ORF77) contains its own TSS (position 93369, Supplementary Table S3) and shares an abundant 3’ end with the upstream *ycf2* transcript at position 93750, probably defined by an RBP (Supplementary Table S5). *ycf15* is a short ORF downstream of *ycf2* whose functionality as a protein-coding gene has been widely debated (97–100), however the presence of discrete 5’ and 3’ ends would be consistent with functionality.

### Terminome-Seq corroborates the production of smRNA footprints from post-transcriptional processing

Despite their high number, TSS only account for a small fraction of 5’ end diversity. Instead, most 5’ and 3’ ends are the result of a maturation and winnowing process where RBPs and secondary structures selectively protect RNAs from degradation by RNases with low sequence specificity (101, 102). Many of these RBPs are members of plant-specific or plant-amplified helical repeat protein families (103), most prominently the PPR family (104). The correlation between smRNA footprints and transcript termini is a key argument which posits that target sequences and secondary structures largely exist to define the ends of functional transcripts (67, 72, 105). It follows that the presence of termini correlating with an smRNA represents an effective way to distinguish true footprints from other smRNAs that are more scattered or whose termini are much more ragged.

A good example of using Terminome-Seq to distinguish smRNA footprints is the *psbB* transcriptional unit (Figure 4), which gives rise to eight accumulating smRNAs (67). These include footprints for three RBPs: HCF107, HCF152 and CRP1; two derived from stem-loops downstream of *psbT*/*psbN* and *petD*, and three smRNAs complementary to the *petD* intron. All but the three intron antisense smRNAs, which may be too unstable to be found in the longer transcripts we sequenced, correlate with termini (Table 1). Another example is the recently-described PPR10-mediated protection of the maize *psaI* 3’ end (73), which correlates with a smRNA from an analogous position in Arabidopsis (genomic position 59475, see Supplementary Figure S6 and Supplementary Table S5). Across the transcriptome, we were able to identify (using the 10 RPM threshold) 45 5’ ends and 44 3’ ends which correlate with the termini of 81 smRNAs (8 footprints correlate with both 5’ and 3’ ends, 37 with 5’ ends only and 36 with 3’ ends only), accounting for 33.5% of the 242 smRNAs previously described (67, 72). Because of the way in which our libraries were made, these ends belong to the longer transcripts from which the smRNAs were ultimately derived.

The ability to look for *bona fide* footprints, along with the elucidation of the PPR code explaining their binding specificity (106–109), will allow the prediction and discovery of new PPR proteins involved in chloroplast RNA stability. Not all of the hundreds of chloroplast PPRs generate accumulating footprints, as demonstrated by PGR3, which is required for *rpl14* 3’ end formation (73).

According to the footprint model, a single RBP can protect the 5’ and 3’ ends of overlapping transcripts. The original example of this phenomenon is maize PPR10, which in binding the *atpI-atpH* intergenic region protects the *atpI* 3’ and *atpH* 5’ ends (71, 110). Although Terminome-Seq could identify the expected *atpH* 5’ end, only a minor 3’ end mapped to the PPR10-dependent *atpI* 3’ end (Supplementary Figure S6). The major Terminome-Seq 3’ end is located further downstream, inside the *atpH* coding region and does not correlate with a smRNA. Systematic mapping of the *atp* operon termini already revealed that transcripts with overlapping ends are a minority (53) and co-immunoprecipitation showed that PPR10 is preferentially associated with processed *atpH* mRNA rather than processed *atpI* mRNA (67). At a genome-wide level, only ∼10 footprints (including PPR38 and HCF152) can be linked to simultaneous stabilization of 5’ and 3’ ends, suggesting that this is the exception rather than the rule. Secondary structures, on the other hand, can stabilize the 3’ ends of two adjacent transcripts, for example in the cases of *psbT-psbN, petD-rpoA* and *petA-psbJ* (Figure 4B and Supplementary Figure S5). Currently, the degradation pathway of RBP footprints is unknown. Evidence derived from Terminome-Seq suggests is consistent with that PNPase participating in this mechanism, as there are more termini correlating with smRNAs in the PNPase mutant than in WT (42.5% vs. 33.5% of all termini; Supplementary Table S5), however a direct analysis of smRNAs in both genotypes would be required to substantiate this possibility.

### Terminome-Seq of the PNPase mutant points to its potential use in reverse genetics

mRNA processing up to the border of the region protected by RBPs or secondary structures is principally performed by three enzymes, RNase J for 5’ ends and PNPase and RNase II for 3’ ends (77, 79, 110). As expected, Terminome-Seq of the PNPase mutant showed a more pronounced effect on 3’ vs. 5’ termini, although many termini of both types were affected (Figure 5 and Figure 6A). We could confirm that PNPase is required to remove tRNA precursor 5’ extensions subsequent to RNase P processing (Supplementary Figures S7,(22)). Similarly, we confirmed that the 3’ extensions observed in the PNPase mutant (22, 34, 74) correlate with 3’ ends distal to secondary structures or RBP binding sites (Supplementary Figures S5, S6 and S8), two elements known to stall PNPase (110, 111).

Using *pnp1-1* as a proof of concept opens the door to understanding the roles of other chloroplast gene regulators using Terminome-Seq. Concerning 5’ termini, this might include the roles of the sigma factors that specify PEP initiation sites (112) and other PEP-associated proteins (PAPs; 113), a closer view at RNase J specificity, or a better understanding of some of the numerous factors that have been implicated in rRNA maturation (6). Terminome-Seq would be an excellent choice to gain additional insight into poorly-characterized RNases whose targets have only begun to be described (114–116).

### Transcription termination analysis with Terminome-Seq

Transcription termination in plastids has recently been reviewed (117). The endosymbiont hypothesis would predict that chloroplast termination would resemble that of bacteria, which use both Rho-dependent and Rho-independent mechanisms. Rho-independent termination occurs in AT-rich sequences downstream of GC-rich stem-loops, however *in vitro* and *in vivo* assays found that PEP terminates inefficiently at chloroplast stem-loops, although it does recognize certain bacterial terminators. This is in keeping with the longstanding hypothesis that chloroplast stem-loops are involved in transcript stability rather than transcription termination (3), a finding well illustrated by the correlation between 3’ ends and secondary structures (Figure 4B and Supplementary Figure S5). On the other hand, Rho-dependent termination, which works through destabilization of the elongating polymerase (118), would have to rely on an alternative cofactor since Rho is not found in organelles. Such a factor was recently identified by its partial similarity to Rho, RHON1, which assists in transcription termination downstream of *rbcL* (119), binding the transcript between two termini identified here (Figure 6C). The 3’ end in the PNPase mutant is further downstream, suggesting a model similar to *E. coli* where Rho factors initiate termination and exoribonucleases like PNPase produce the precise, mature 3’ ends directly downstream of stem-loops or RBP sites (120). Similarly, MTERF6, a member of a gene family related to a human mitochondrial transcription termination factor, acts in the *trnI* and *petD-rpoA* regions (65, 117, 121). As additional factors involved in termination are discovered, Terminome-Seq offers a tool to decipher their sites of action.

### Conclusion

Several high-throughput RNA-Seq-based strategies have recently been used to gain an unprecedented genome-level understanding of chloroplast RNA biology. It is now possible to study RNA abundance, splicing and editing (26–28), to monitor translation rates with ribosome profiling (24, 122), and to infer protein binding sites from smRNA sequencing (67, 72, 105). Terminome-Seq now creates access to a single-nucleotide resolution map of the full set of chloroplast RNA termini. We anticipate that this strategy, combined with the other plastome-wide approaches, will be instrumental in deciphering mechanisms such as transcription (from initiation to termination), as well as the roles of RNases and RBPs in shaping the chloroplast transcriptome.

