## Supplementary Figure for "Systematic sequencing of chloroplast transcript termini from *Arabidopsis thaliana* reveals >200 transcription initiation sites and the extensive imprints of RNA-binding proteins and secondary structures"

#### SUPPLEMENTARY FIGURE LEGENDS

**Supplementary Figure S1: Terminome-Seq strategy**

Total leaf RNA was used to create three different RNA-Seq libraries. For 5' end analysis, optional TAP treatment was completed and a 5' adapter was ligated, followed by fragmentation and ligation of a 3' adapter. For 3' end analysis, the 3' adapter was ligated first. The first adapter ligation prior to fragmentation ensures that all cDNAs amplified for library sequencing represent an end naturally present in the RNA population containing either a 5'-P or 3'-OH. Technical details are in the Materials and Methods section.

**Supplementary Figure S2: Reproducibility between replicates for A) WT and B) *pnp1-1* Terminome-Seq**

Replicates of the indicated library samples were graphed based on RPM for a given end. Correlation coefficients for each strand are given to the bottom left of each graph.

**Supplementary Figure S3: Identifying and validating TSS using Terminome-Seq**

**A)** TSS within *ndhF* were mapped using 5' RACE**.** The overlapping *ycf1.1* and *ndhF* gene models are shown with Terminome-Seq results for the minus strand below (see key for color scheme). Black arrowheads represent the 3' primers used for 5' RACE, for which the corresponding stained gels of the PCR reactions are shown.

**B)** Comparison between +TAP coverage (red) vs –TAP coverage (blue) for selected 5' ends. The derived ratio used to determine TSS status is indicated above the graph. Genomic positions 53093 and 74411 have been described as P*atpE* -430 (Kapoor *et al.*, 1994; Ghulam *et al.*, 2013) and P*psbN* -32 (Zghidi *et al.*, 2007), respectively, but do not reach the $\frac{+\mathrm{TAP}}{-\mathrm{TAP}}$>10 threshold. Genomic position 32525 previously described as P*psbD* -186 (Hanaoka *et al.*, 2003; Hoffer and Christopher, 1997) is likely to be a processed end. Genomic position 122076, previously described as the *ndhA* -66 processed 5' end (Zoschke *et al.*, 2016), gives a strong indication of being a TSS.

**Supplementary Figure S4: Terminome-Seq coverage of the *atpI/H/F/A* (A) and *ndhH/A/I/G/E/psaC/ndhD* (B) gene clusters**

The corresponding gene models are shown below, with exons in gray and introns in white. –TAP 5' ends are in blue and 3' ends are in green; bent arrows represent TSS inferred from +TAP data. Numbered peaks and promoters refer to features listed in Supplementary Table S4.

**Supplementary Figure S5: Transcript termini at secondary structures**

Terminome-Seq coverage around predicted secondary structures in the WT and PNPase mutant. Gene models are represented as open arrows and color coding of ends is provided in an inset. Genome positions where ends accumulate are indicated with black arrows. The stem-loops matching described smRNA are highlighted in green, and an RBP footprint is in blue.

**Supplementary Figure S6: Transcript termini surrounding known and putative RBP sites**

Terminome-Seq coverage near binding sites for HCF152, PPR10, HCF107, CRP1, HCF145, RAP, SOT1, PGR3, PPR5, PPR38, EMB175 and CRR2 is indicated for WT and the PNPase mutant (*pnp1-1*). Gene models are represented as open arrows and color coding of ends is provided in an inset. Genome positions where ends accumulate are indicated with black arrows. RBP footprints matching described smRNAs are highlighted in blue.

**Supplementary Figure S7: Transcript termini upstream of tRNAs**
Comparison of Terminome-Seq coverage for WT and *pnp1-1* in selected tRNA-encoding regions. Gene models are represented as open arrows and color coding of ends is provided in an inset. Genome positions where ends accumulate are indicated with black arrows, and smRNA locations are highlighted in blue. TSS for tRNAs are indicated by bent arrows. Additional data, for *trnG*, is shown in Supplementary Figure S8.

**Supplementary Figure S8: Transcript termini in intergenic regions**
Comparison of Terminome-Seq coverage for WT and *pnp1-1* in selected intergenic regions. Gene models are represented as open arrows and color coding of ends is provided in an inset. Genome positions where ends accumulate are indicated with black arrows, and smRNA locations are highlighted in blue.

**Supplementary Figure S9. Terminome-Seq coverage for the rRNA operon**
The rRNA gene model is shown below (grey arrows), excluding the tRNAs. The dashed area highlights the second hidden break in the 23S rRNA at position 106441, and the vertical arrow the 23S 3' extension in *pnp1-1*. Tick marks are every 1,000 nt.

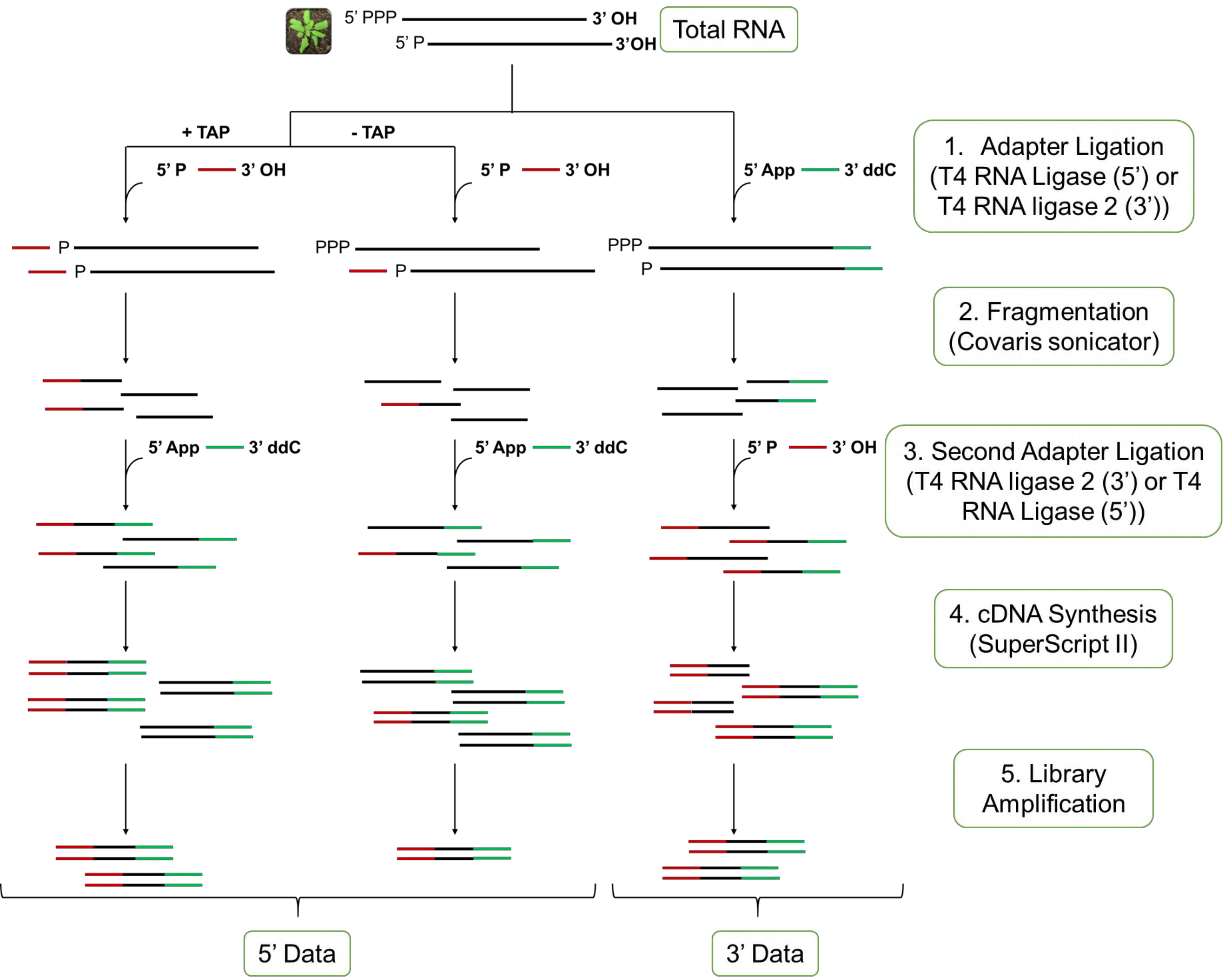


**Supplementary Figure S1**

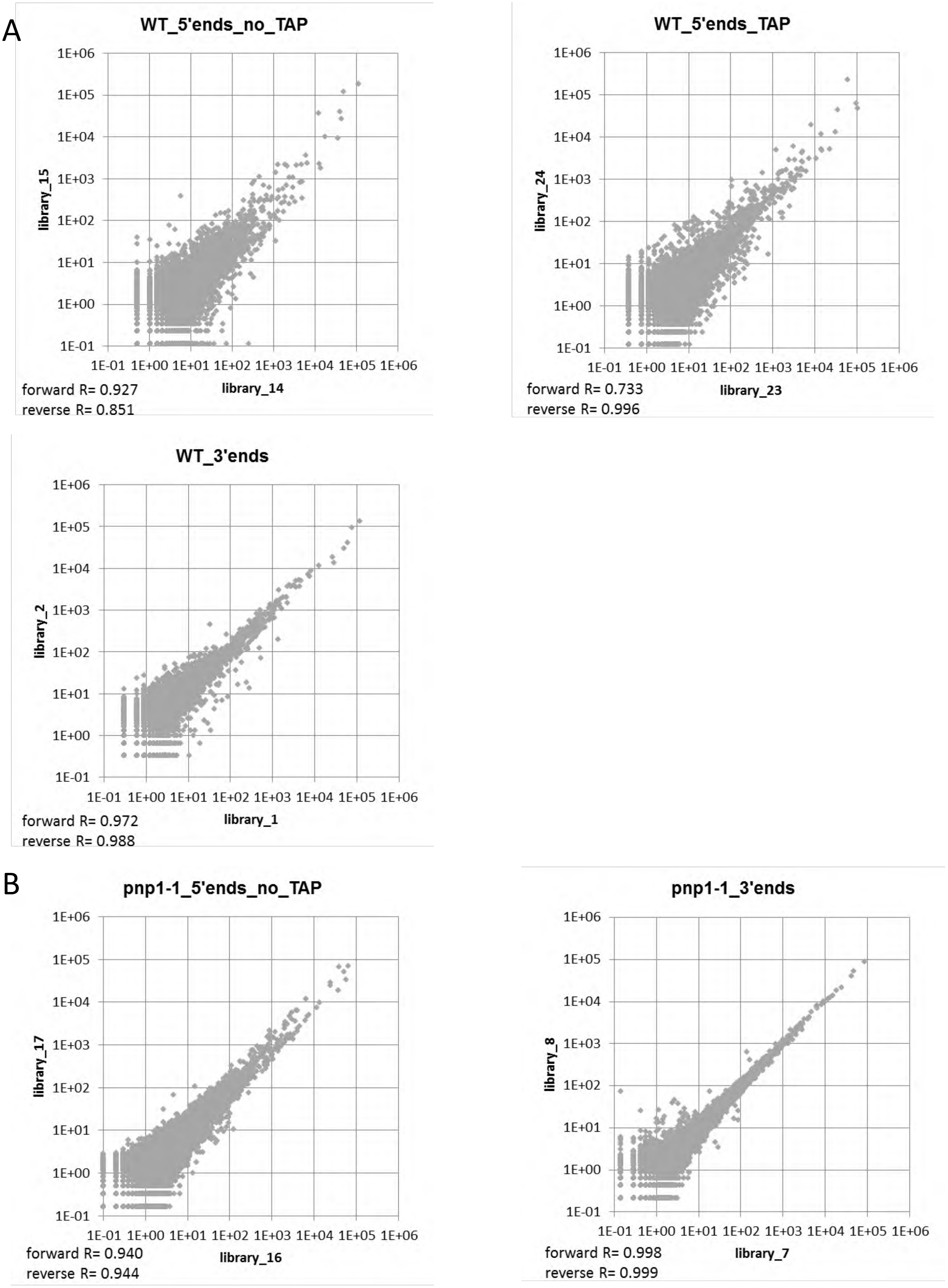


**Supplementary Figure S2**

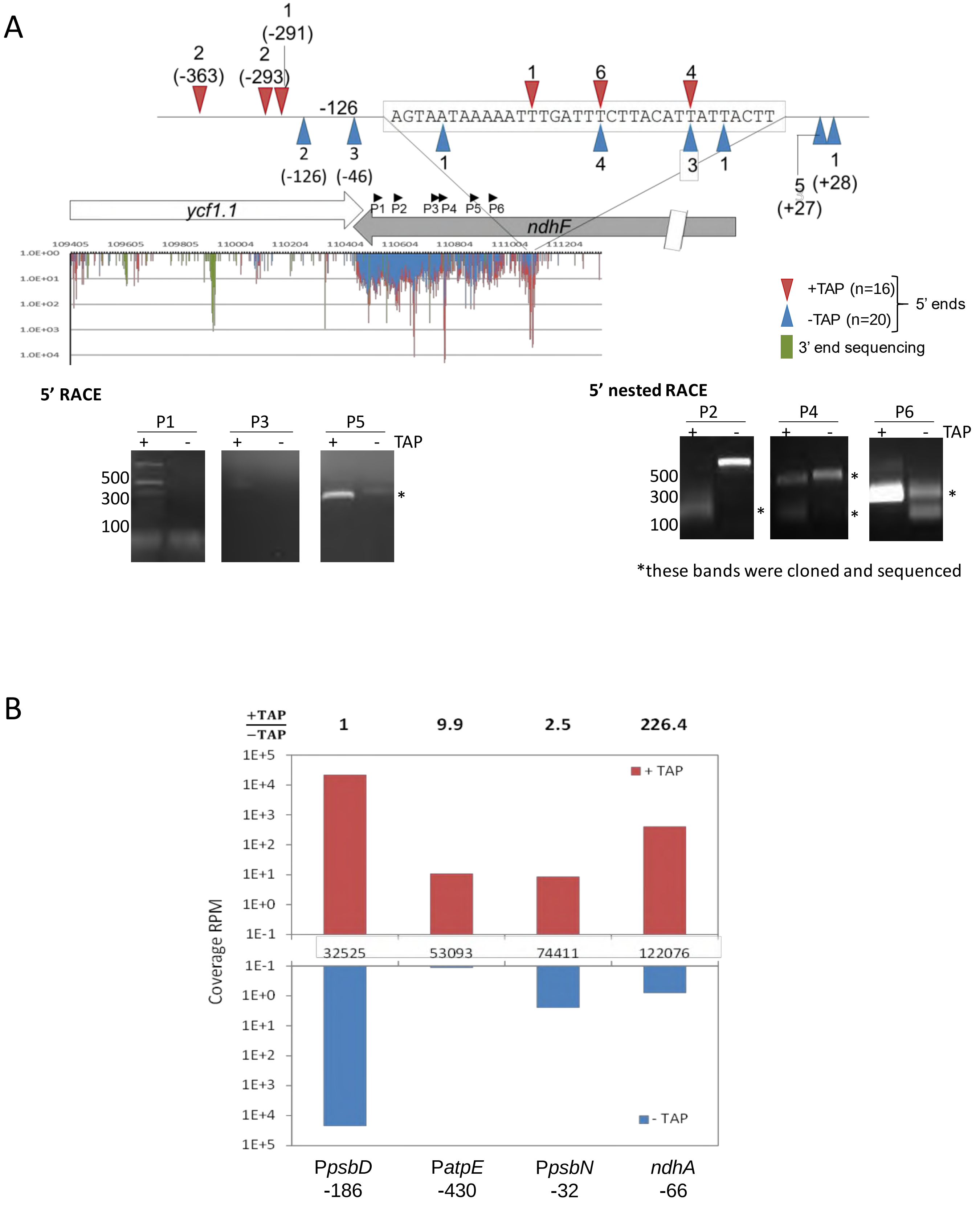


### Supplementary Figure S3

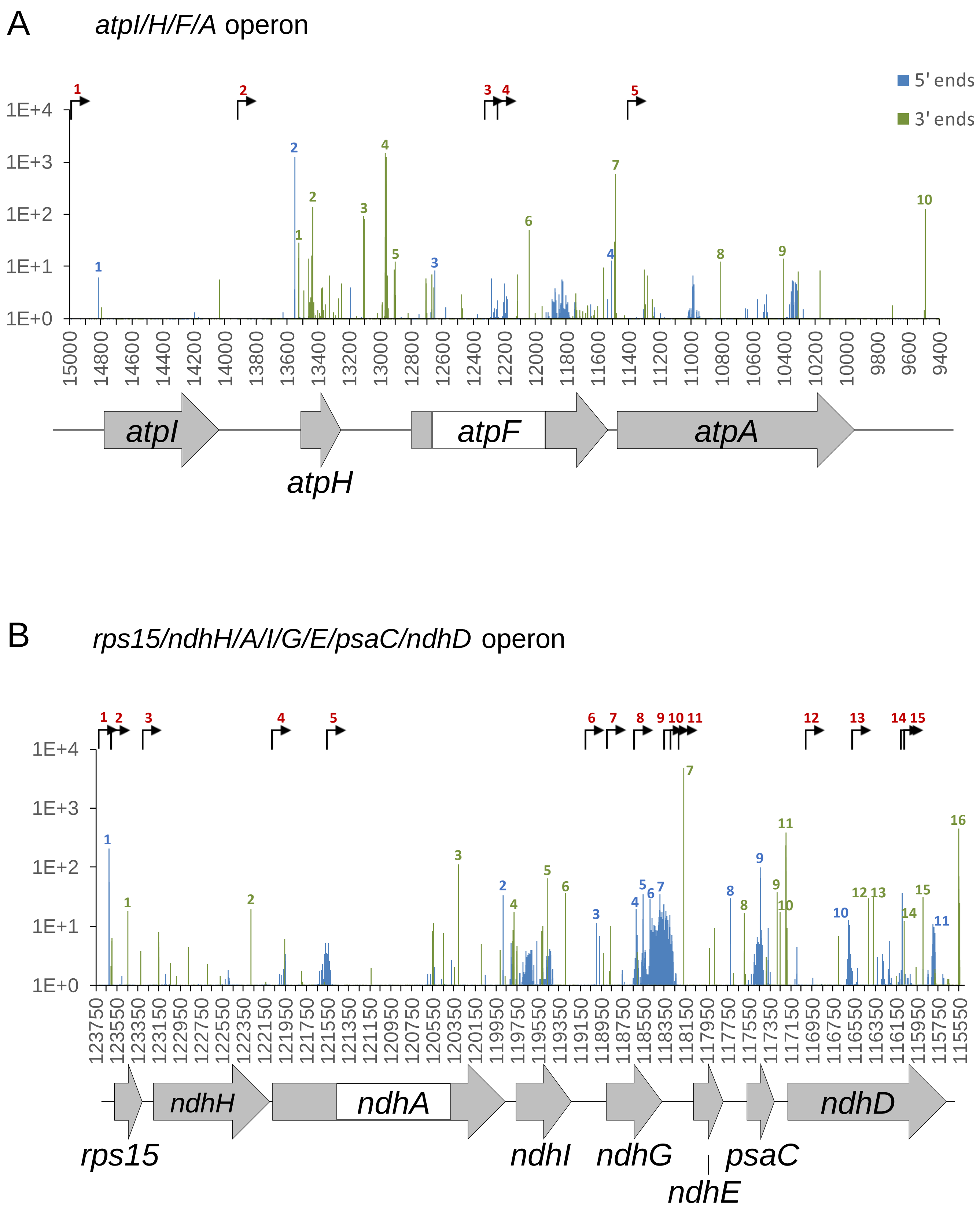


**Supplementary Figure S4**

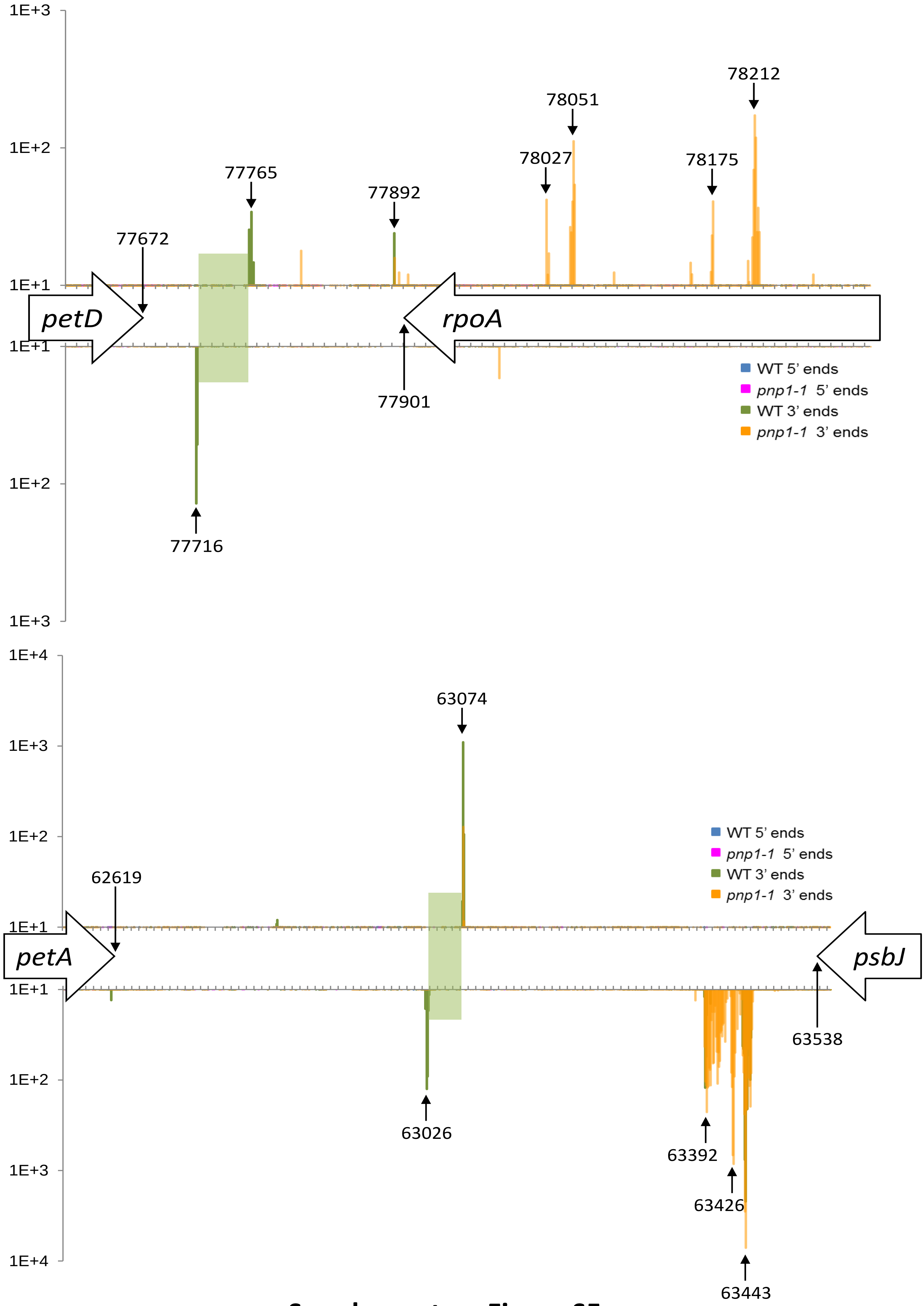


**Supplementary Figure S5**

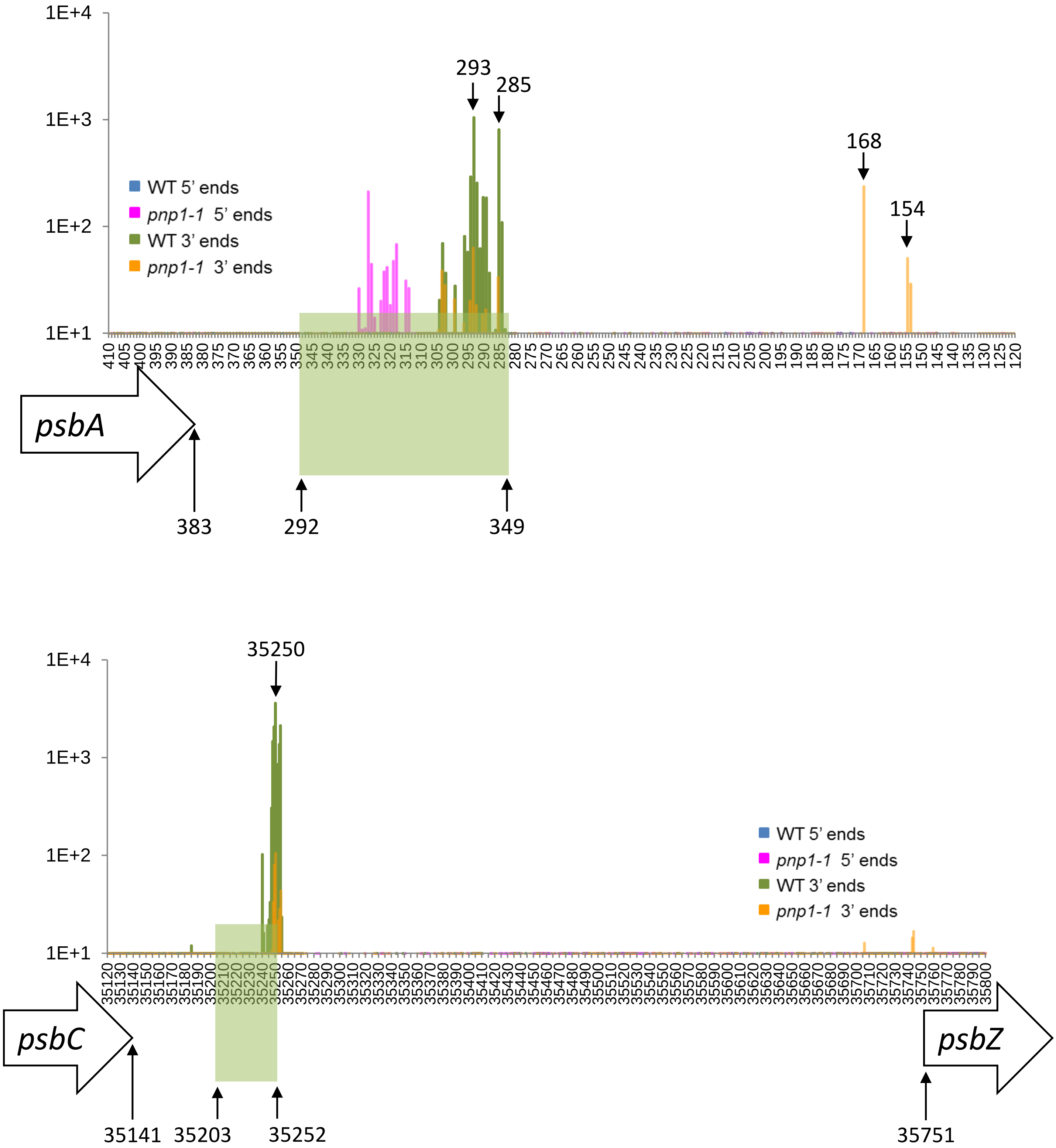


**Supplementary Figure S5**

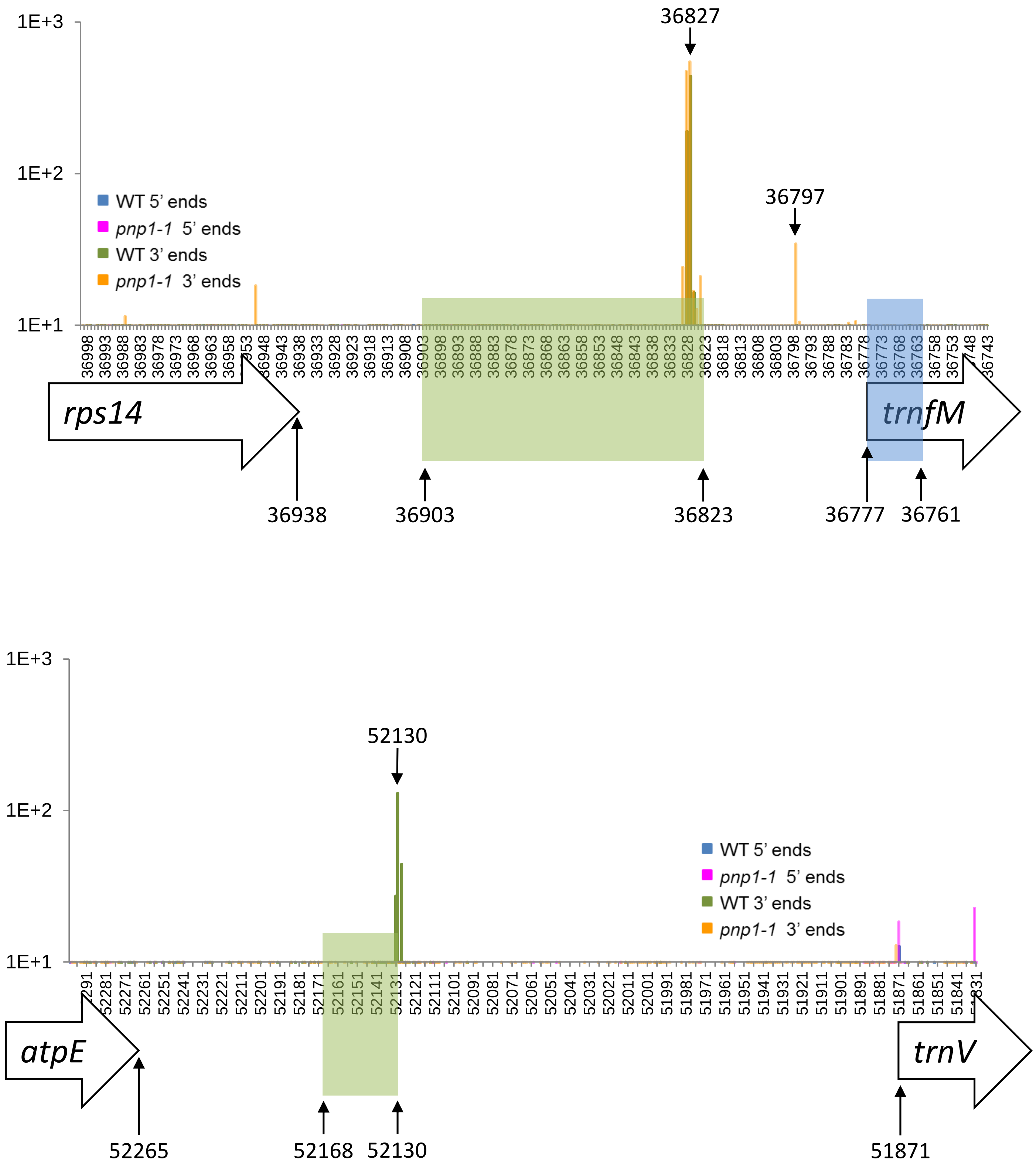


**Supplementary Figure S5**

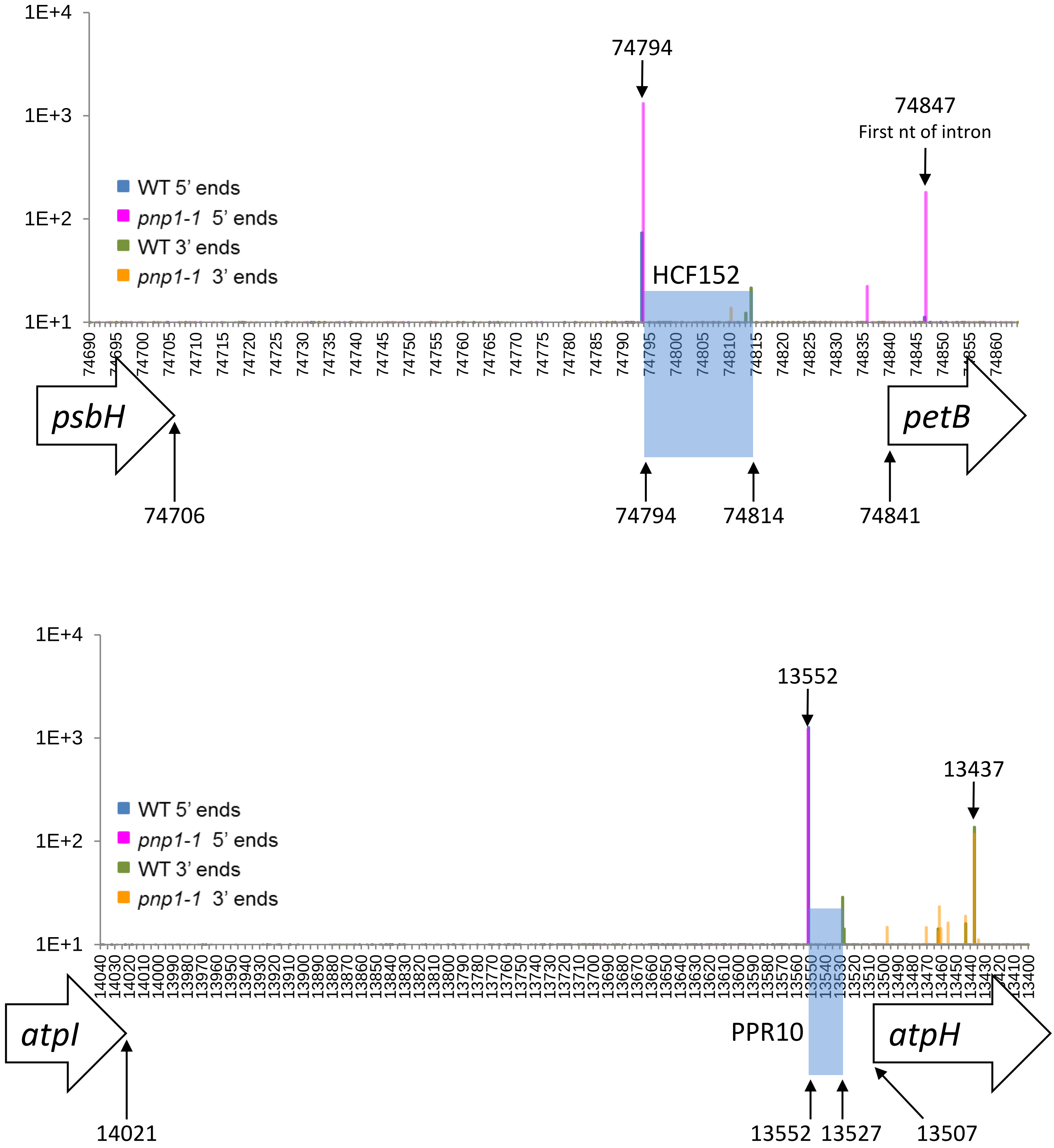


**Supplementary Figure S6**

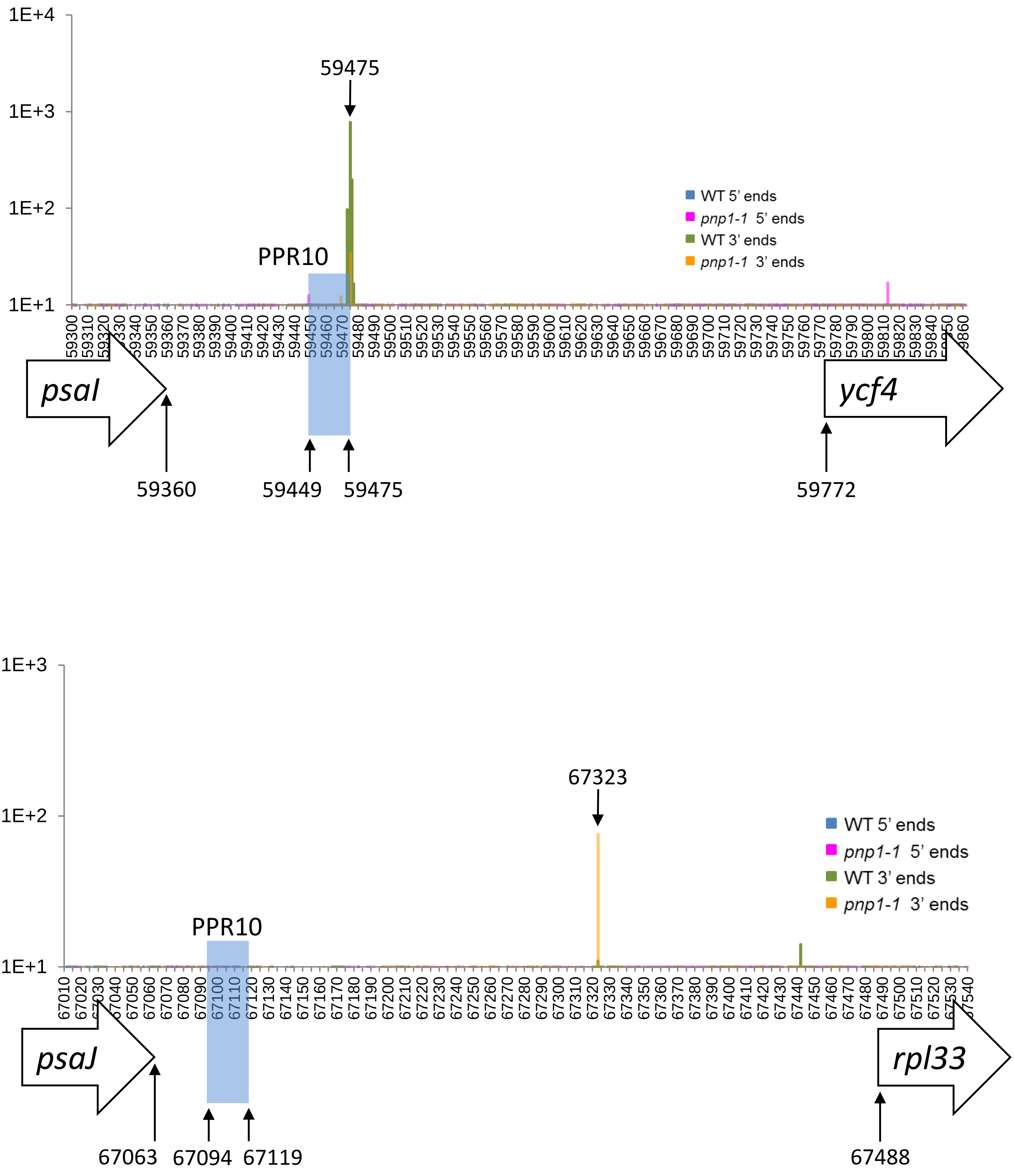


**Supplementary Figure S6**

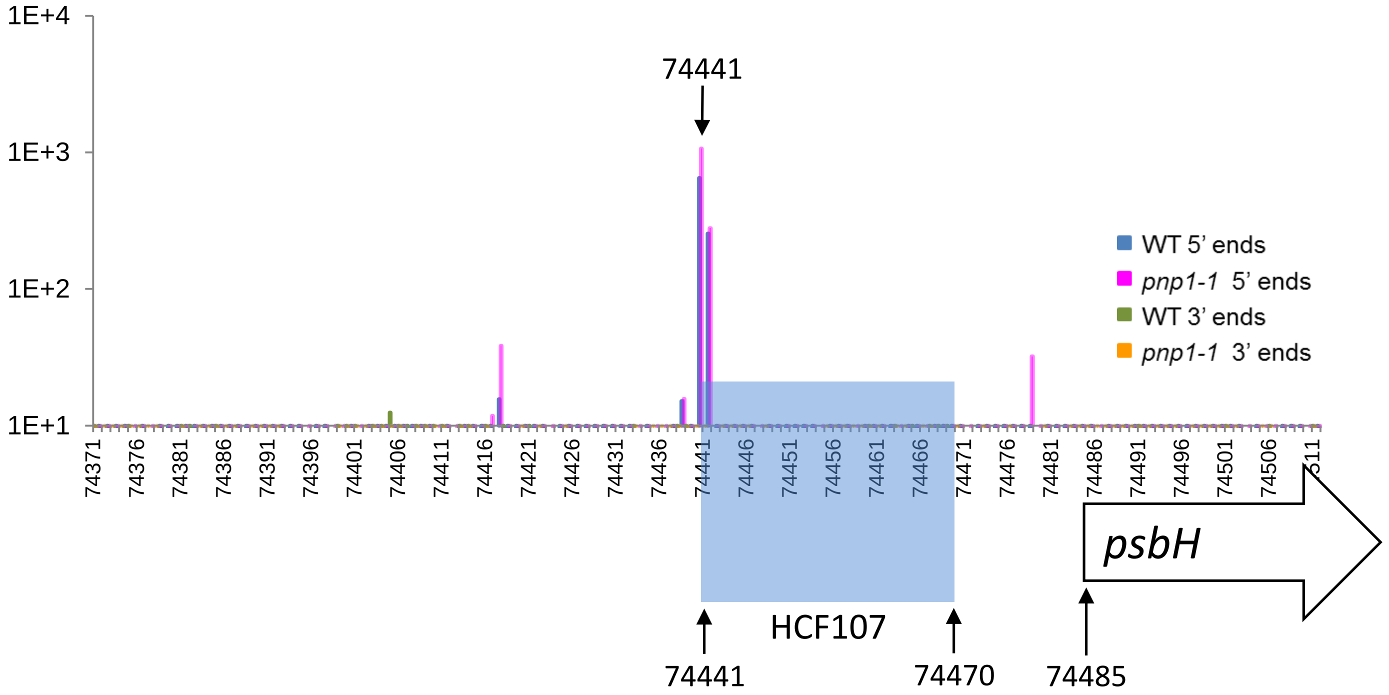
1E+4
1E+3
1E+2
1E+1
74441
WT 5' ends
pnp1-1 5' ends
WT 3' ends
pnp1-1 3' ends
HCF107
psbH
74441
74470
74485

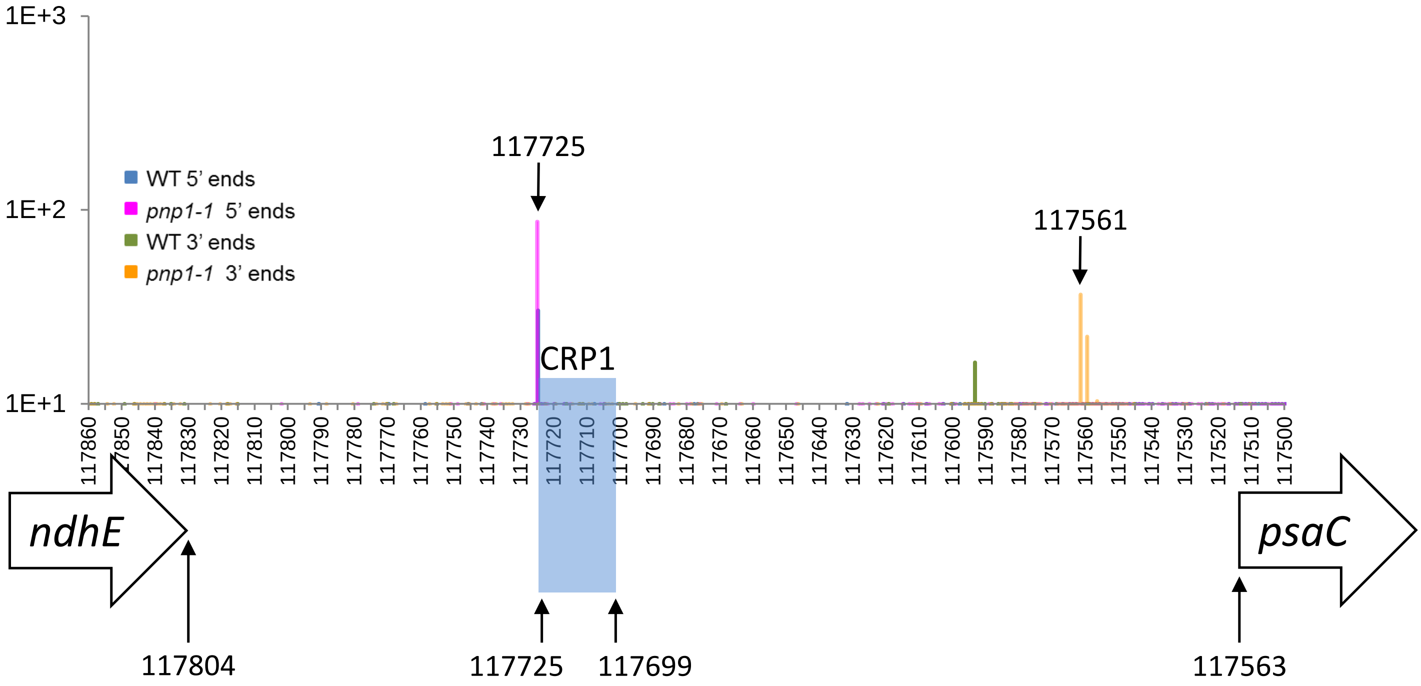
1E+3
1E+2
1E+1
WT 5' ends
pnp1-1 5' ends
WT 3' ends
pnp1-1 3' ends
117725
117561
CRP1
ndhE
psaC
117804
117725
117699
117563

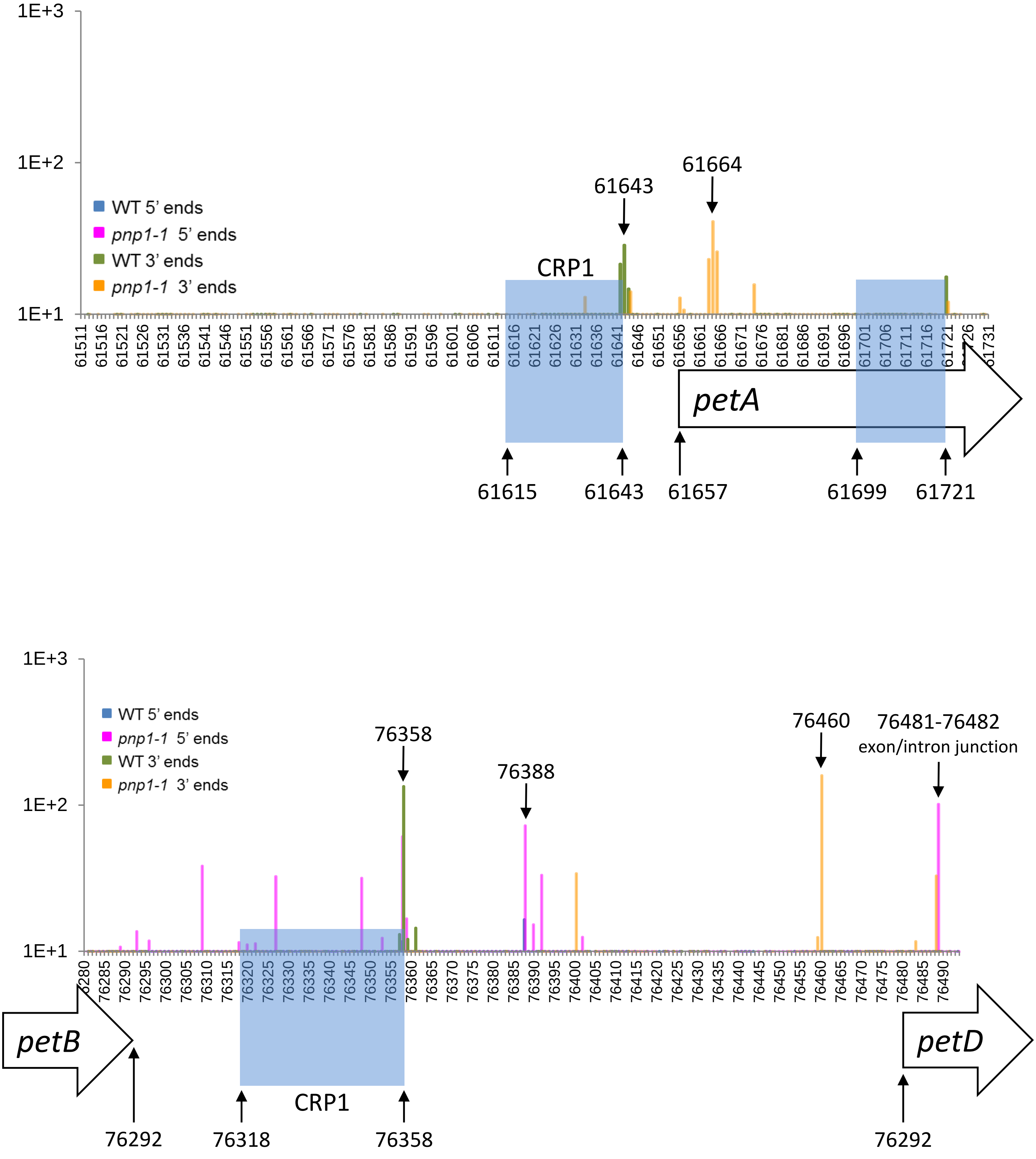


**Supplementary Figure S6**

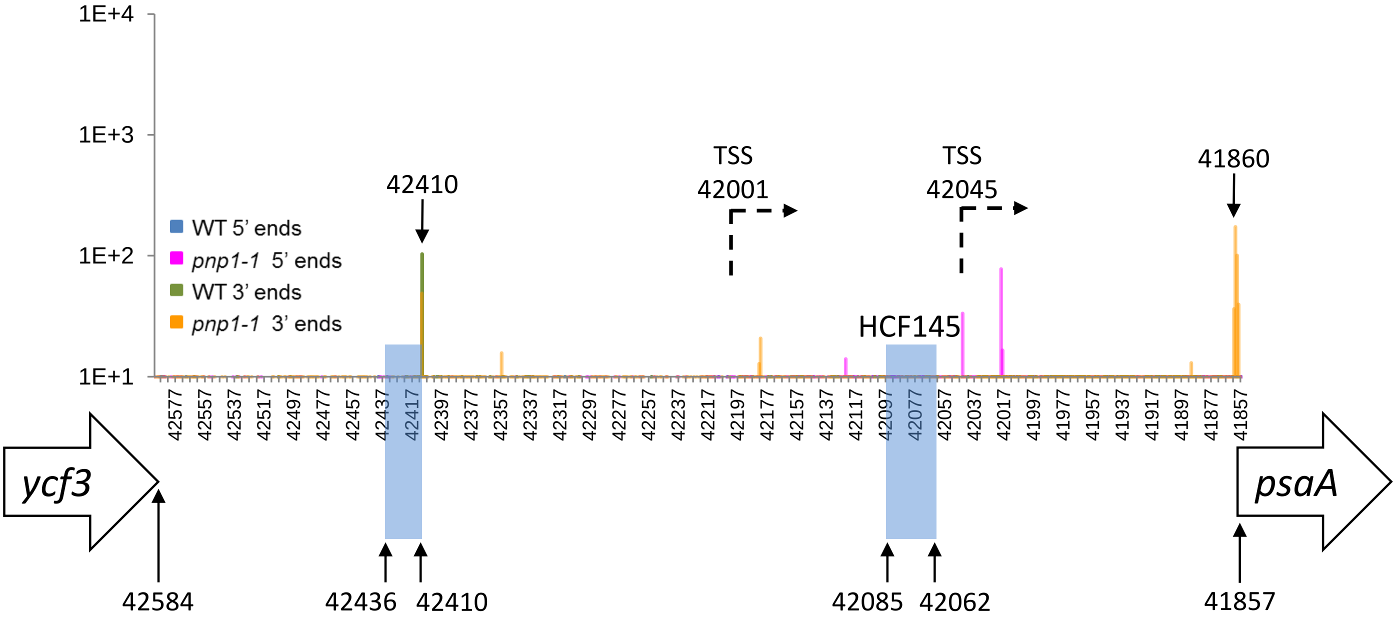


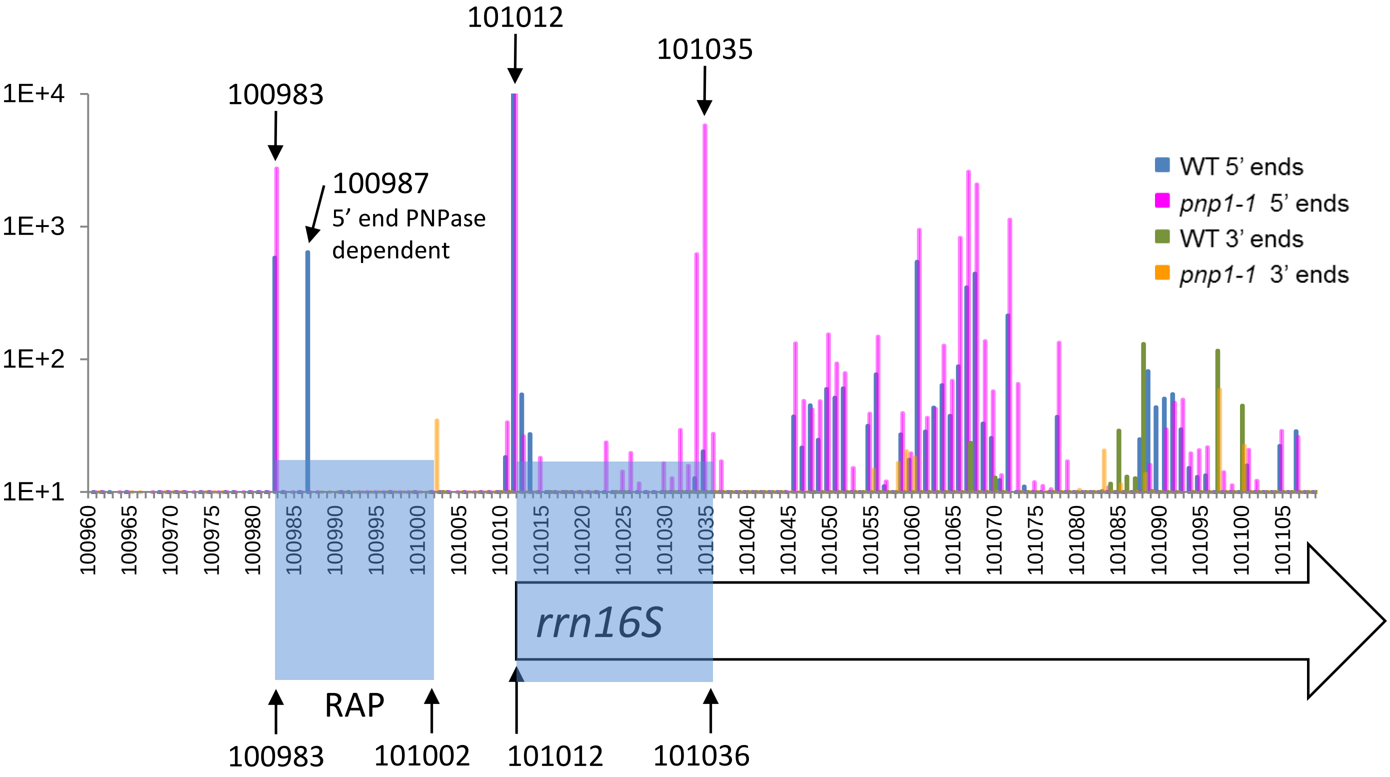


**Supplementary Figure S6**

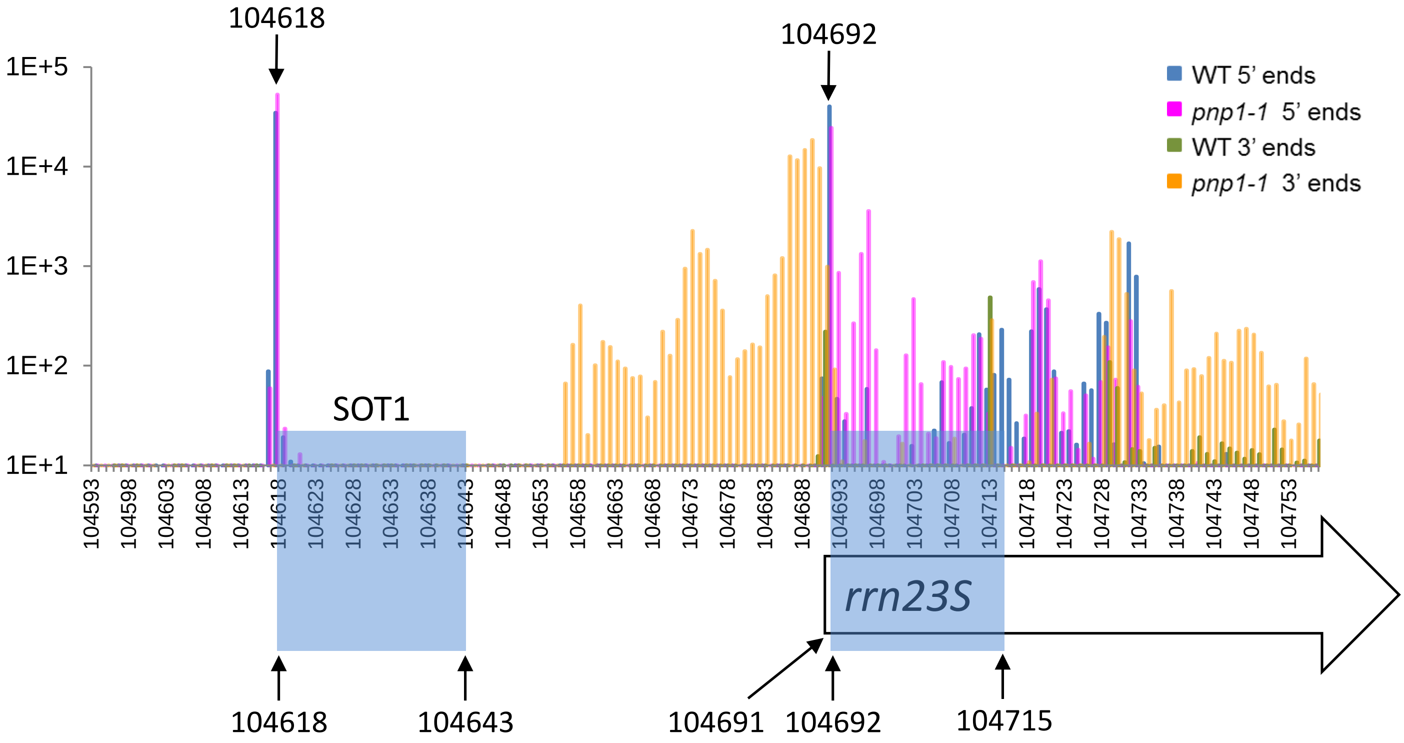


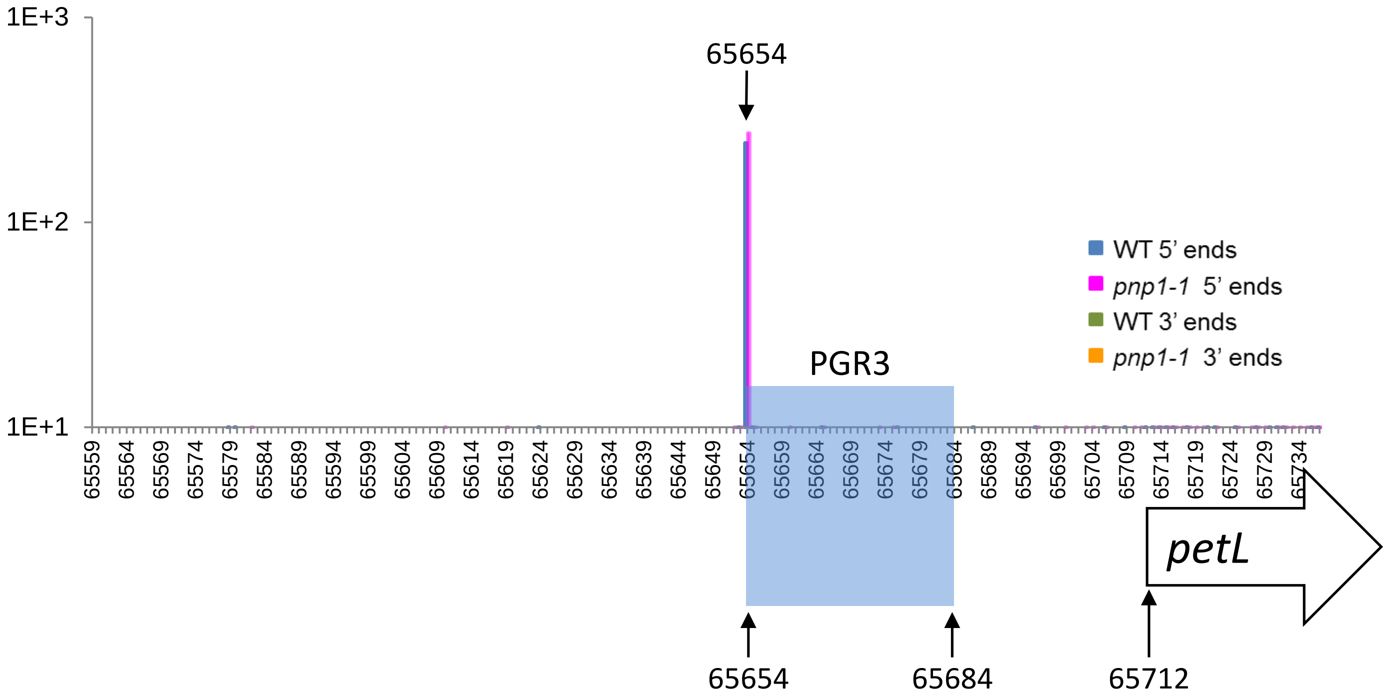


**Supplementary Figure S6**

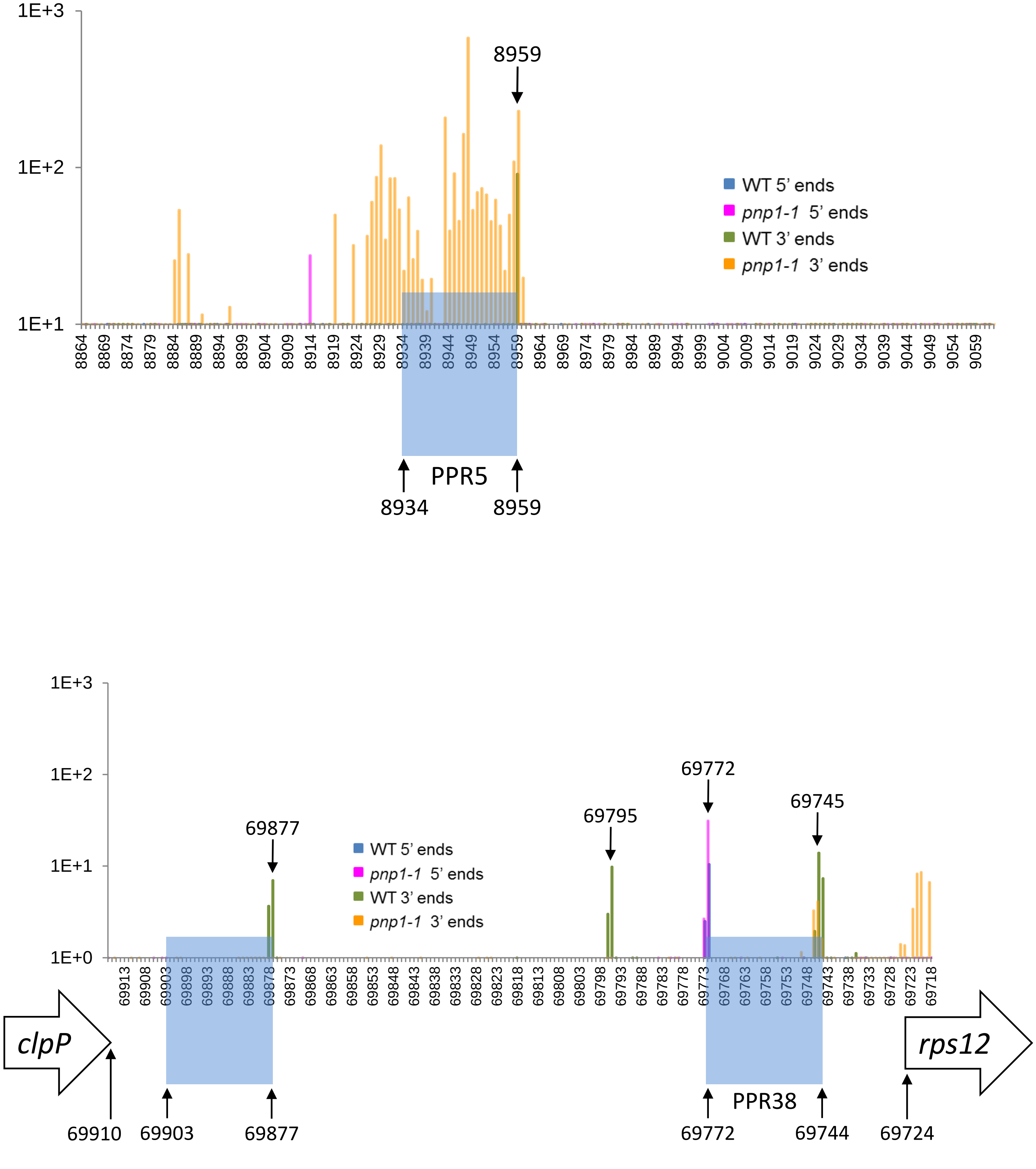


**Supplementary Figure S6**

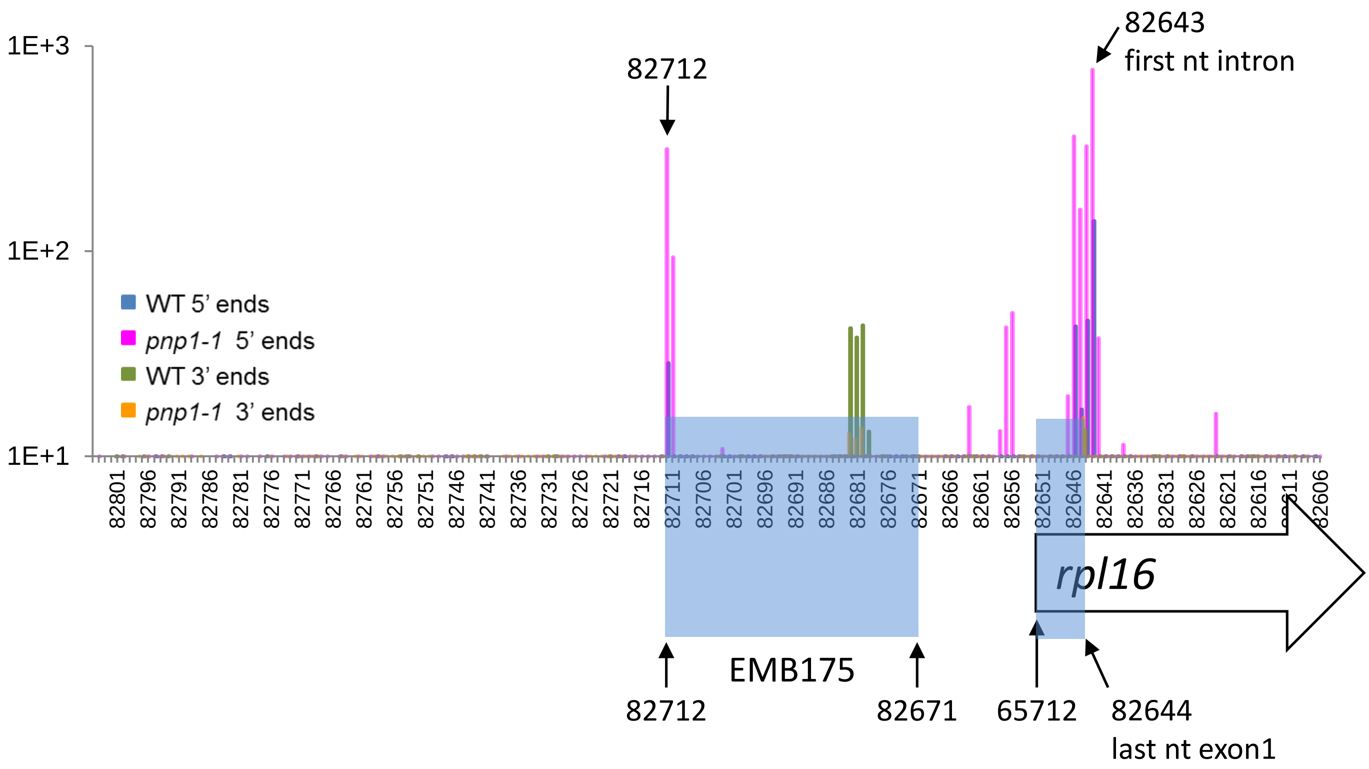


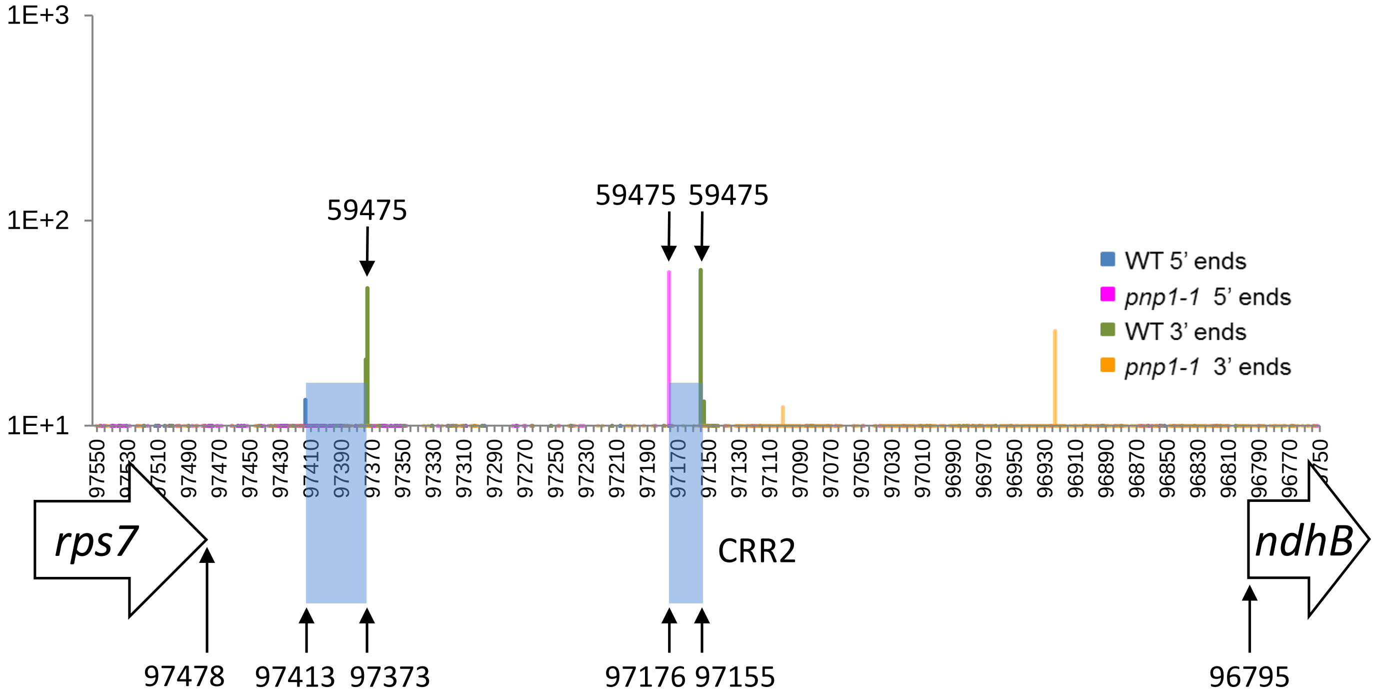


**Supplementary Figure S6**

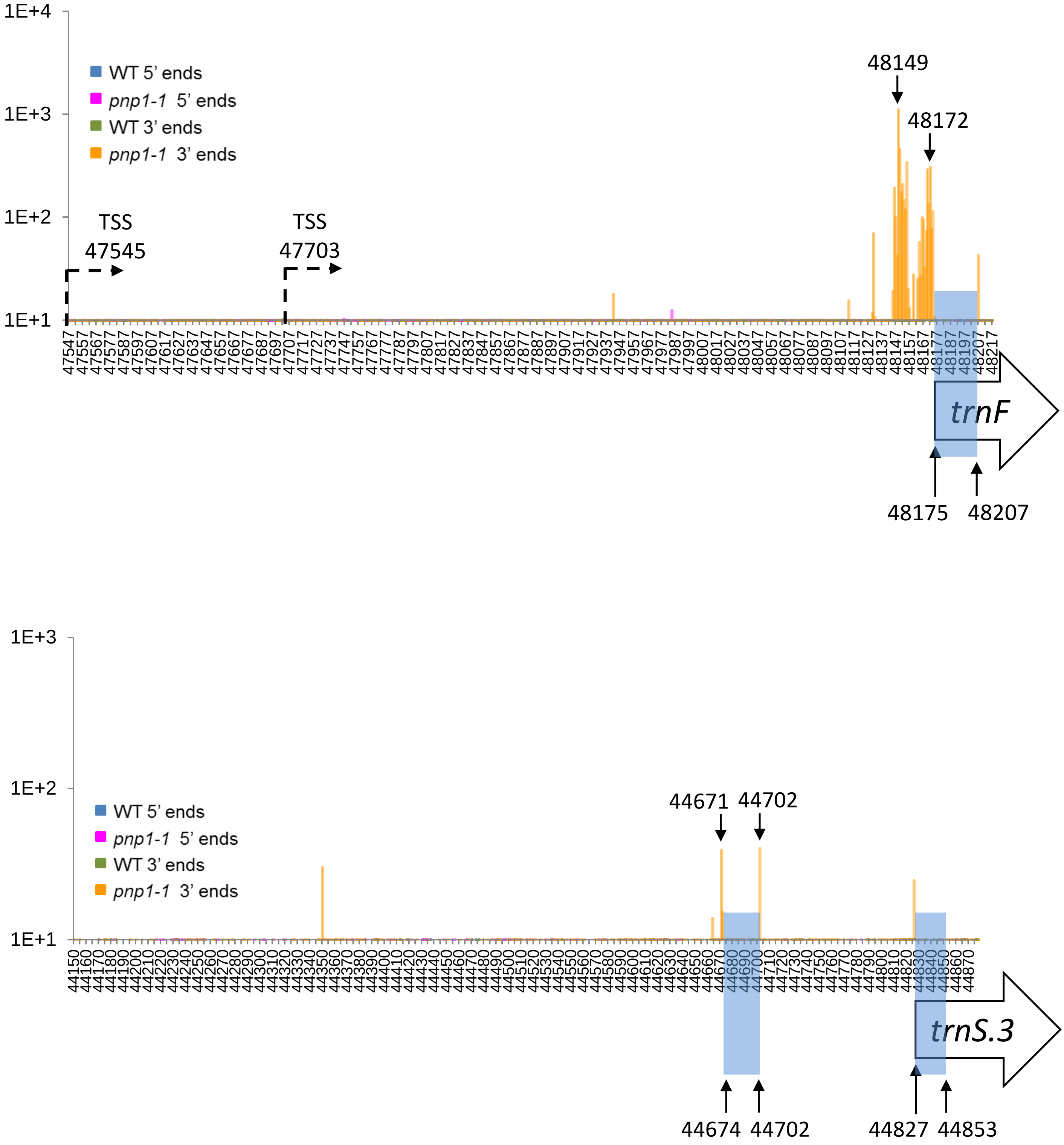


**Supplementary Figure S7**

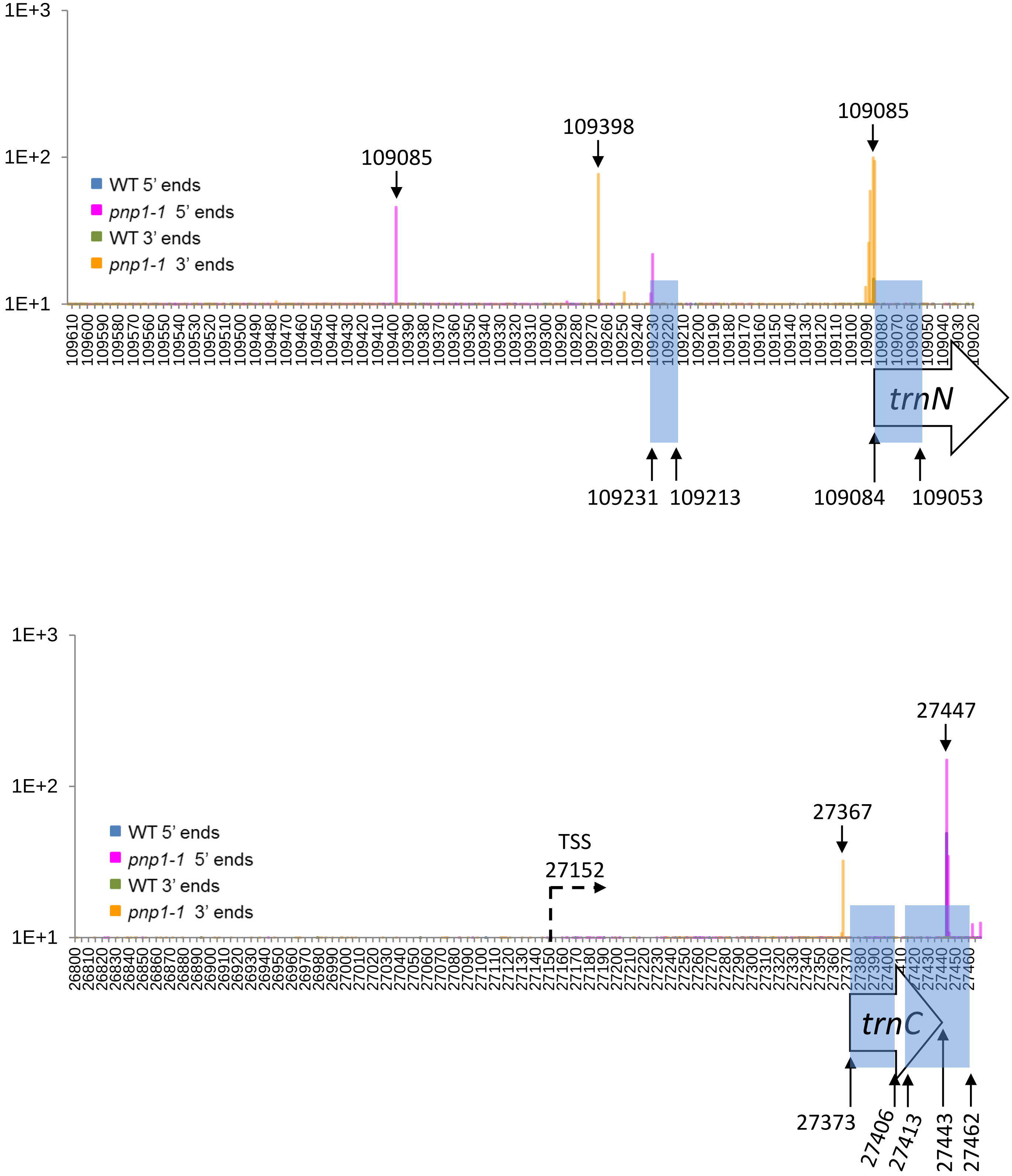


**Supplementary Figure S7**

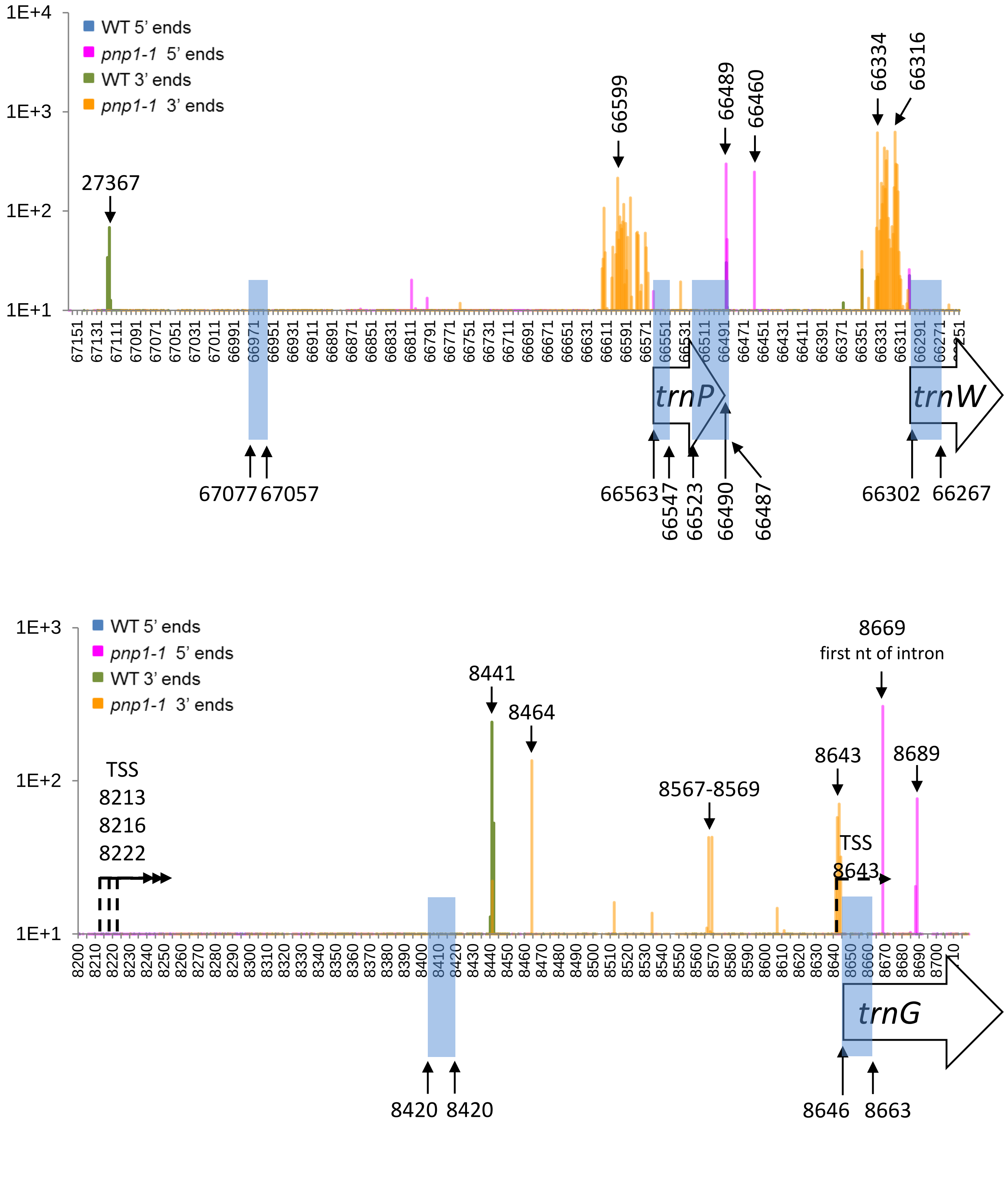


**Supplementary Figure S7**

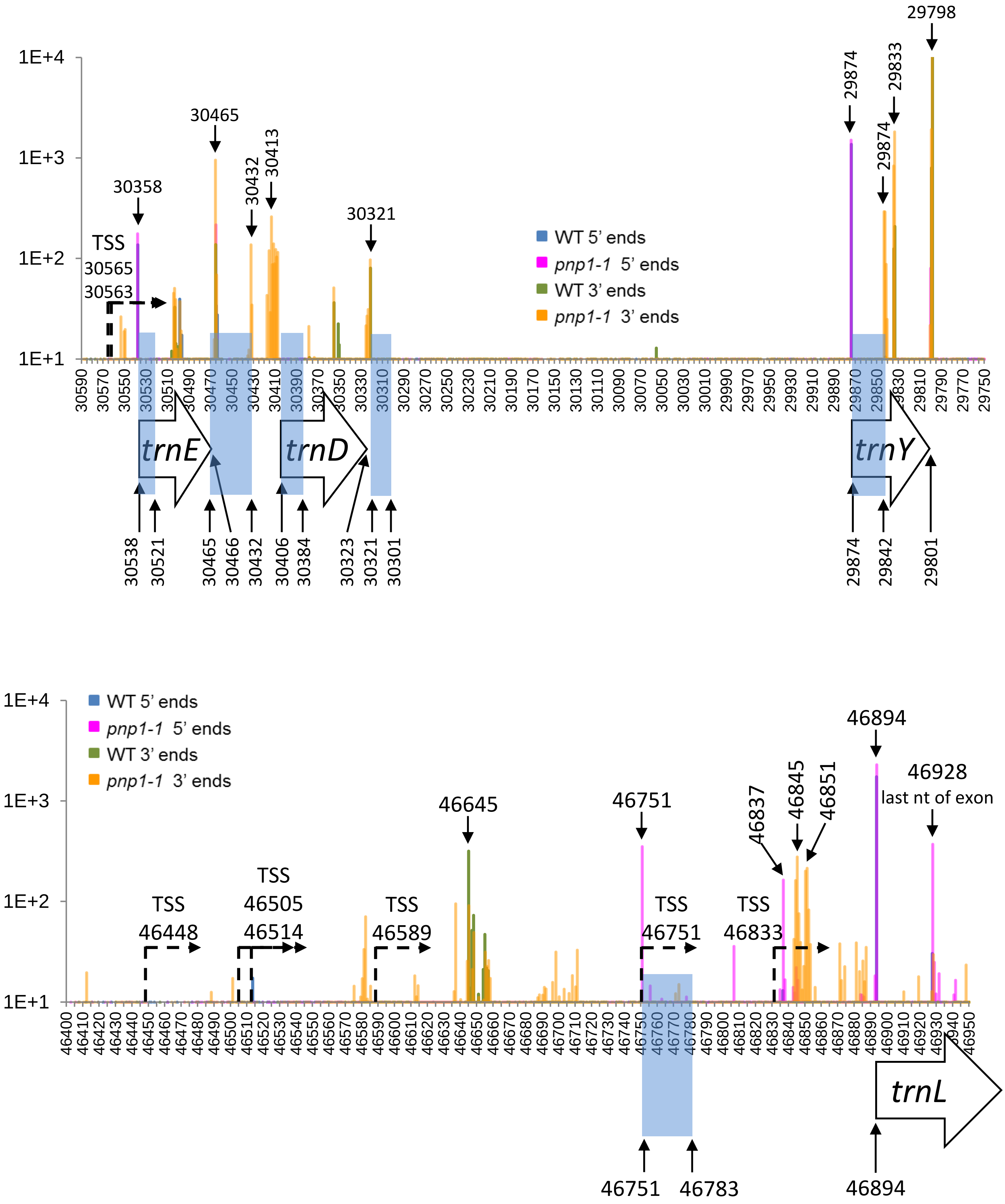


**Supplementary Figure S7**

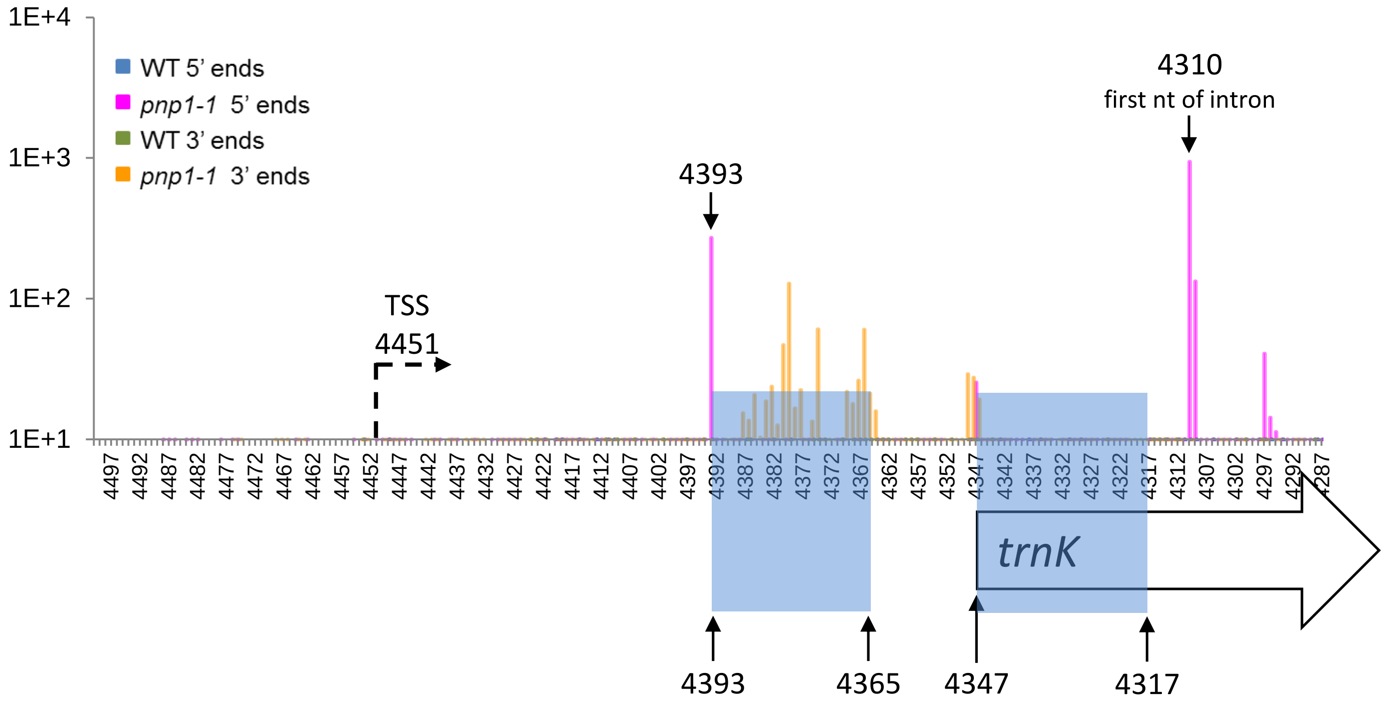


**Supplementary Figure S7**

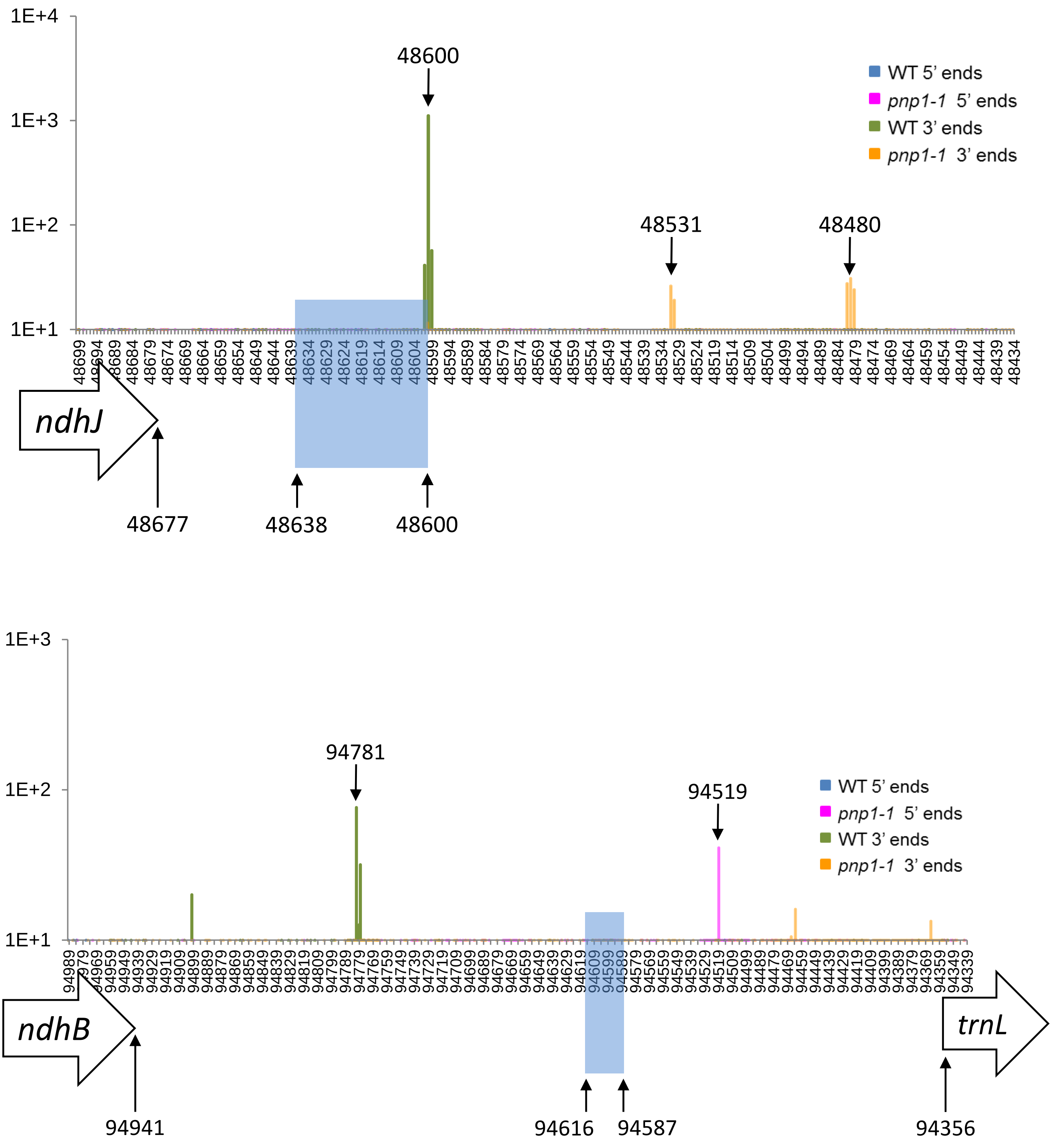


**Supplementary Figure S8**

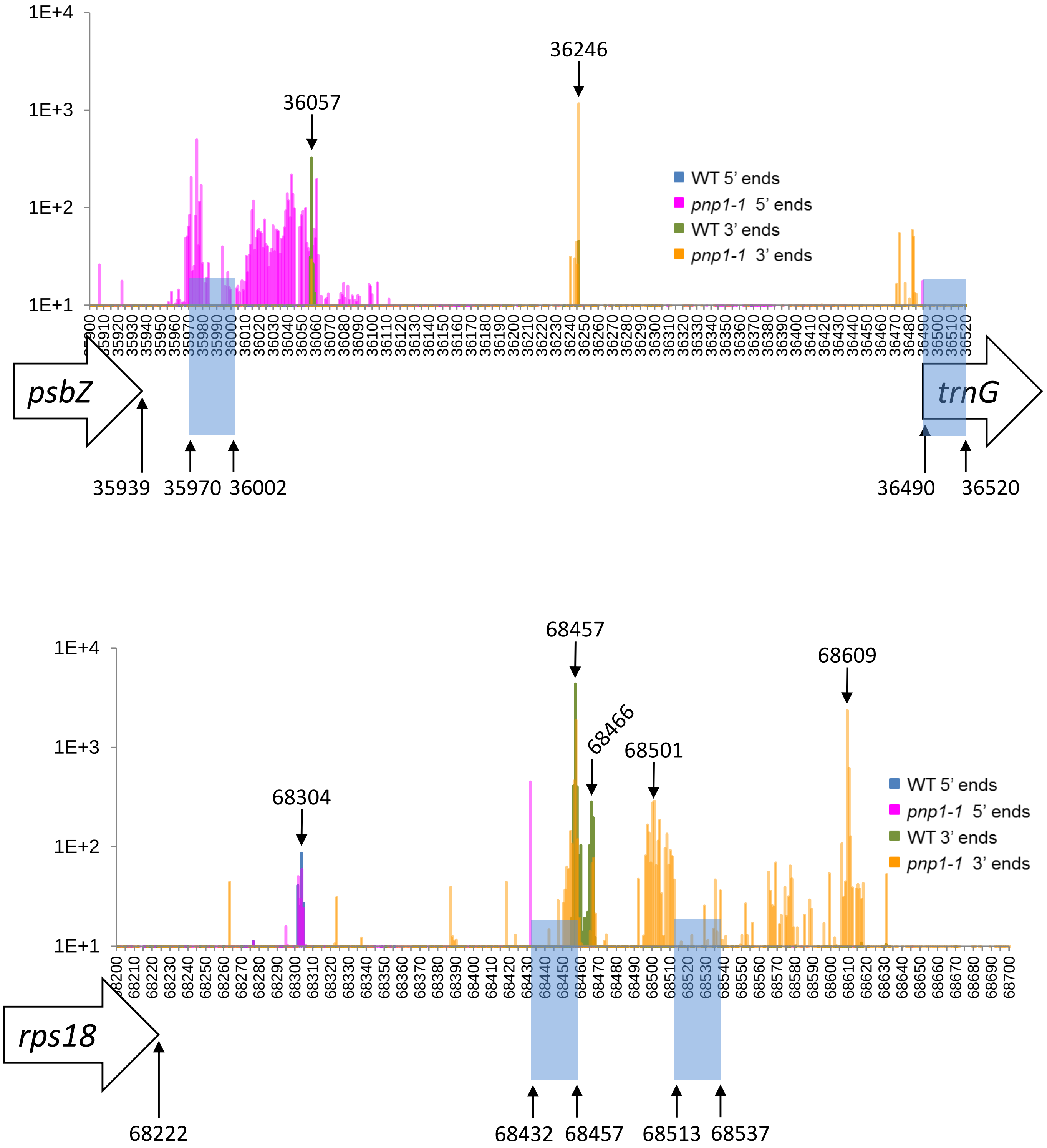


**Supplementary Figure S8**

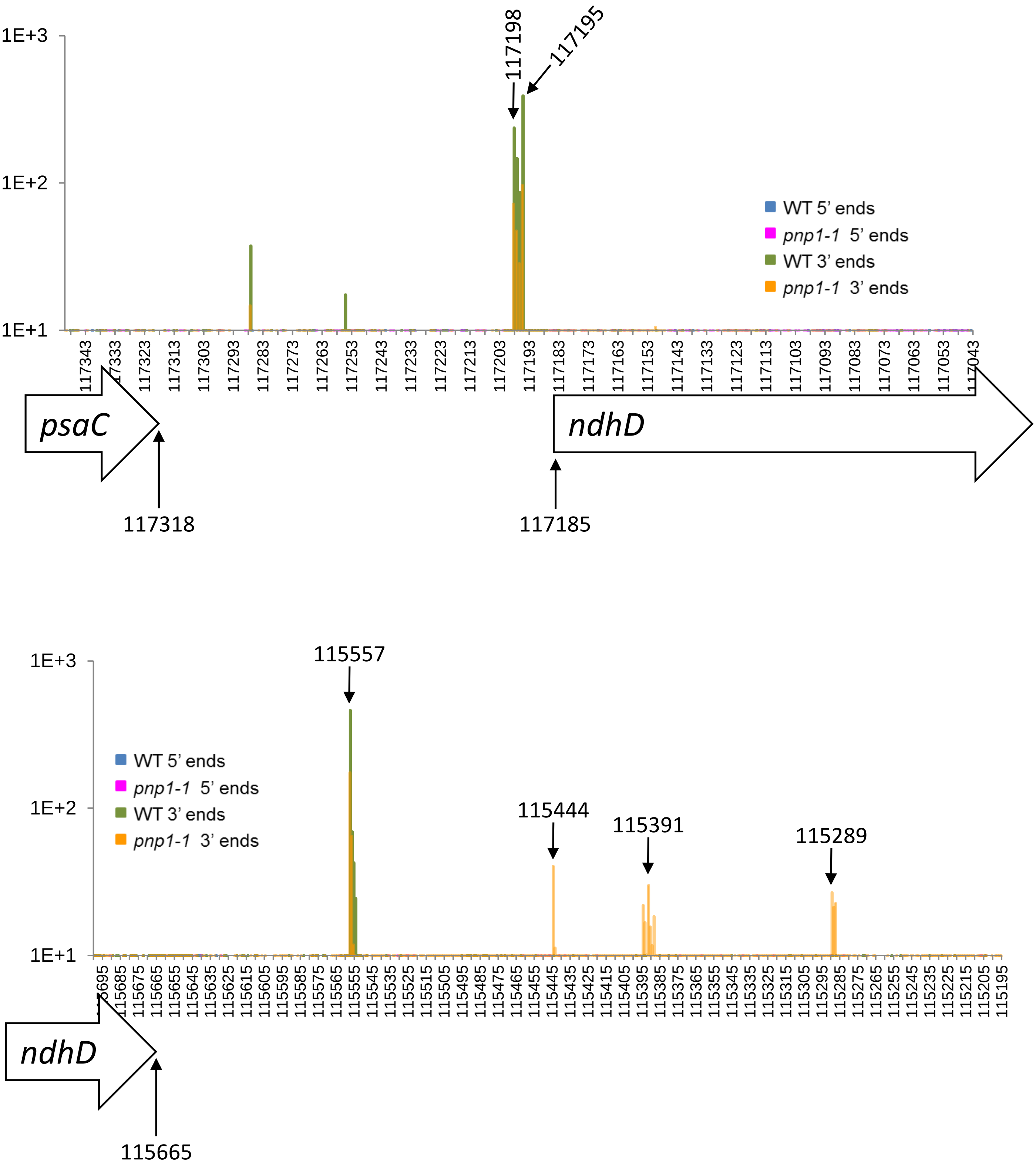


**Supplementary Figure S8**

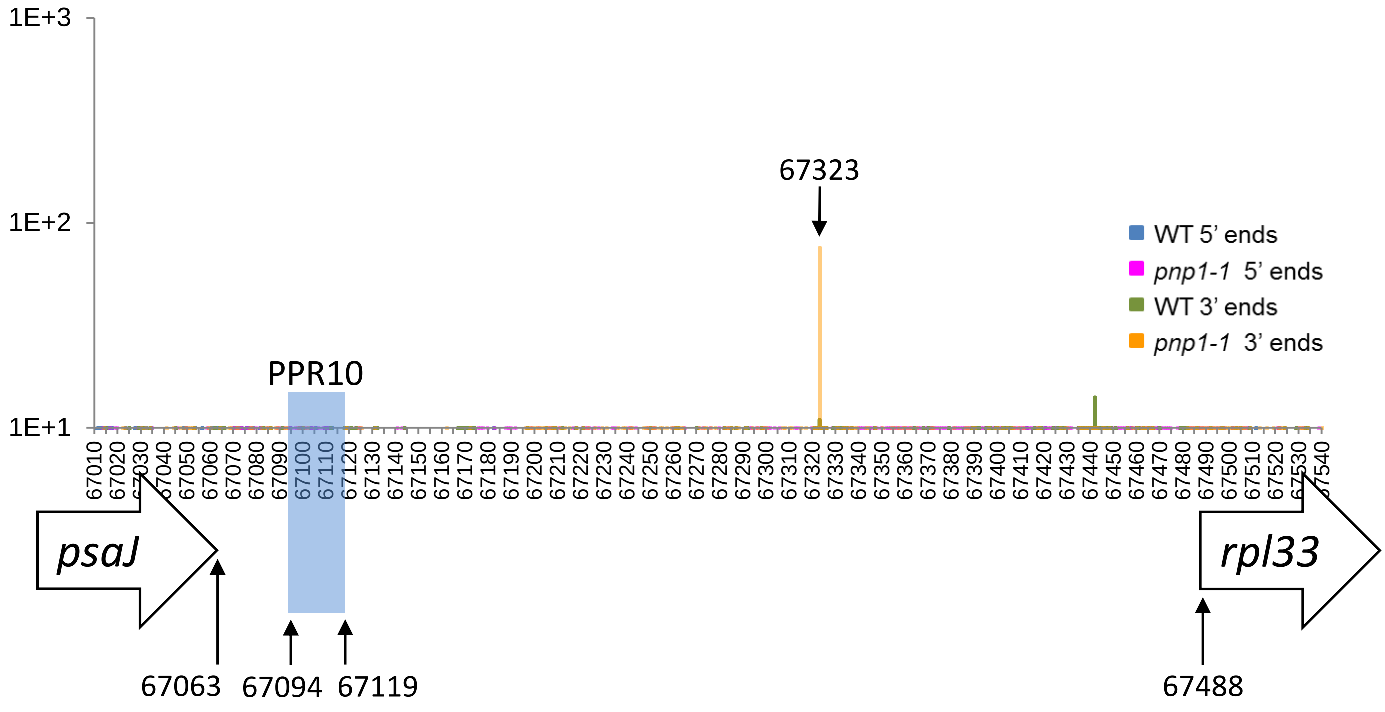


**Supplementary Figure S8**

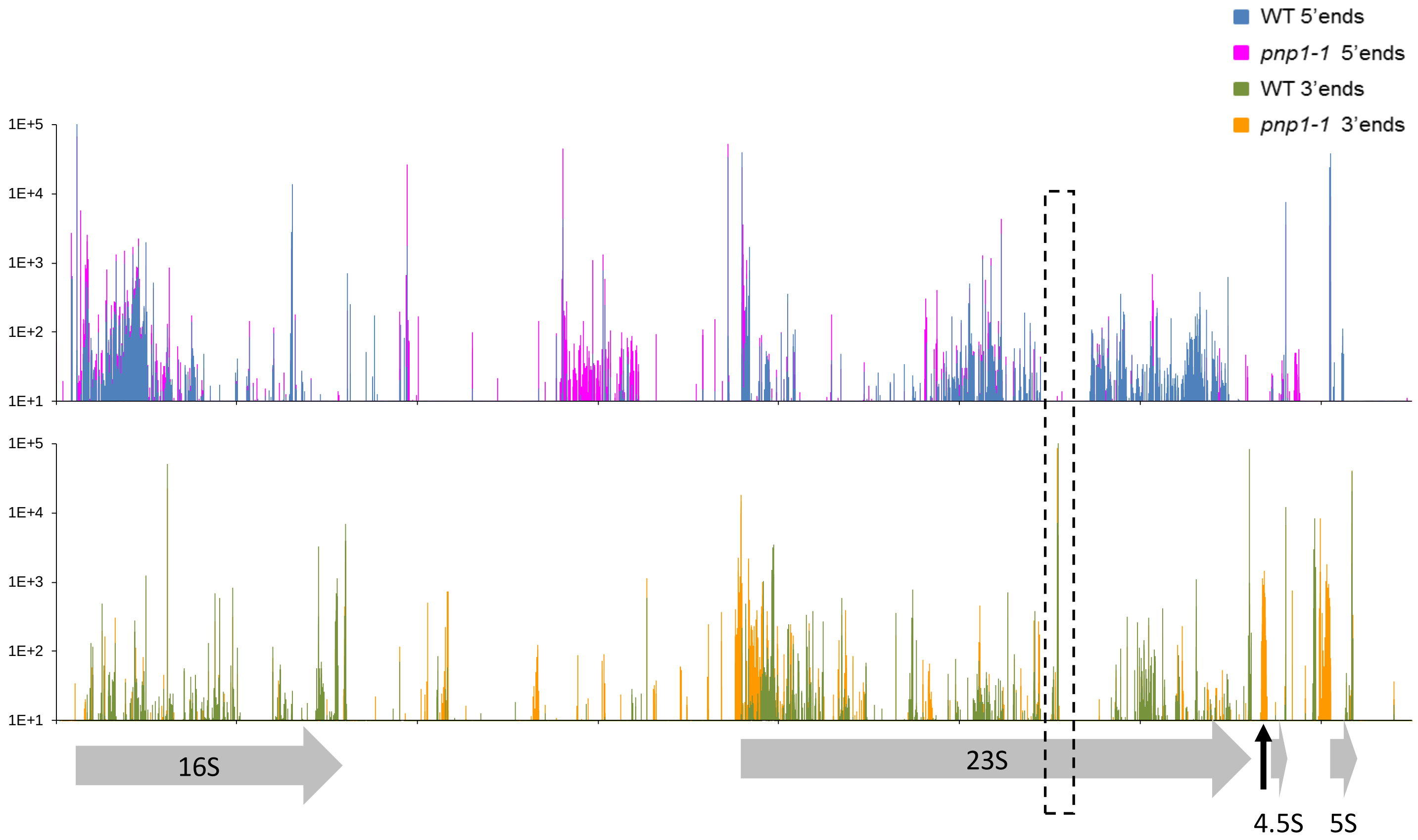


**Supplementary Figure S9. Terminome-Seq coverage for the rRNA operon**
The rRNA gene model is shown below (grey arrows), excluding the tRNAs. The dashed area highlights the second hidden break in the 23S rRNA at position 106441, and the vertical arrow the 23S 3' extension in *pnp1-1*. Tick marks are every 1,000 nt.
